# Divergent CRD-Dependent Mechanisms Govern RAS Isoform-Selective Recruitment of CRAF and ARAF

**DOI:** 10.64898/2026.05.08.723844

**Authors:** Shrhea Banerjee, Sravani Malasani, Shriti Banerjee, Maria Celeste López Vásquez, Shane McSorley, Zhihong Wang

## Abstract

RAF kinases interpret signals from the three major RAS isoforms to initiate MAPK pathway activation, yet the molecular logic that governs isoform-specific RAS recruitment and the early events that relieve RAF autoinhibition are not yet fully understood. In particular, how the modular N-terminal regulatory architecture of CRAF and ARAF, anchored by the multifunctional cysteine-rich domain (CRD), discriminates among HRAS, KRAS, and NRAS has remained a central unresolved question. Here, we combine quantitative biophysical measurements with structural and dynamic analyses to define how RAS isoform identity and CRD engagement shape the earliest steps of RAF activation. These studies reveal unexpectedly divergent modes of RAS recognition between CRAF and ARAF and expose previously unappreciated functions of the CRD in modulating RAS affinity and intramolecular regulatory contacts. We further identify a direct link between RAS binding and destabilization of RAF autoinhibition, providing mechanistic insight into how RAS initiates the transition from an inactive monomer to an activation-competent assembly. Finally, we show that emerging KRAS inhibitors variably perturb KRAS-CRAF interactions, offering insight into how these therapeutics influence early RAS-RAF signaling events. Together, this work uncovers distinct biophysical principles that govern RAS-RAF selectivity and reveals a regulatory role for the CRD that reframes our understanding of RAF activation and its dysregulation in RAS-driven cancers.

**Significance:** Proteins in the RAS-RAF signaling pathway control cell growth and are frequently mutated in cancer. Despite their importance, how different RAS proteins selectively recruit RAF kinases has remained incompletely understood. This study reveals that the cysteine-rich regulatory region of RAF plays a central role in distinguishing RAS isoforms and controlling RAF activation. These insights clarify early steps in MAPK signaling and may guide the development of improved therapies targeting RAS-driven cancers.

## Introduction

The mitogen-activated protein kinase (MAPK) signaling pathway is a highly regulated, three-tier kinase cascade that governs cellular responses to extracellular signals.(Cope et al., 2018; Lavoie & Therrien, 2015) The RAS/RAF/ERK signaling cascade is a principal driver of cell proliferation and survival in response to extracellular mitogenic signals. Signal initiation occurs when GTP-loaded RAS recruits RAF kinases to the plasma membrane, triggering RAF activation and subsequent activation of downstream MEK and ERK proteins.(Cope et al., 2018; Rajakulendran et al., 2009; Vojtek et al., 1993; Yu et al., 2023) Activated ERK in turn phosphorylates a diverse set of nuclear transcription factors to drive cellular responses.

RAF kinases, ARAF, BRAF, and CRAF, are the principal effectors that transmit signals from RAS to the MAPK cascade. RAF kinases share conserved domain organization comprising three regulatory regions (CR1-CR3).(Cope et al., 2018) CR1 contains the Ras-binding domain (RBD) and cysteine-rich domain (CRD), which together mediate effector recognition and phospholipid interactions.(Brtva et al., 1995; Cope et al., 2018; Fang et al., 2020; Hu et al., 1995; Trebino et al., 2023) CR2 contributes to 14-3-3 binding and autoinhibitory stabilization, while CR3 contains the kinase domain and the second 14-3-3 binding site that governs dimerization.(Cutler et al., 1998; Rajakulendran et al., 2009) Within this modular framework, the CRD has emerged as a multifunctional regulatory hub: it coordinates zinc, binds the plasma membrane (Fischer et al., 2007; Li et al., 2018), participates directly in RAS recognition (Cookis & Mattos, 2021; Tran et al., 2021), and forms extensive autoinhibitory contacts with both 14-3-3 and the kinase domain.(Cope et al., 2019; Gunderwala et al., 2019; Martinez Fiesco et al., 2022; Park et al., 2019) In its inactive form, monomeric RAF is maintained in an autoinhibited state through intramolecular interactions, until extracellular signaling triggers release of this autoinhibited state and subsequent dimerization.(Park et al., 2019) RAF activation is tightly controlled by dimerization; with catalytic activity emerging only upon formation of homo- or heterodimers.

Although the RAF isoforms share conserved domain architecture, they differ substantially in intrinsic kinase activity, regulatory constraints, and signaling output.(Baljuls et al., 2007; Chong & Guan, 2003; Cope et al., 2019) BRAF exhibits the highest basal activity, followed by CRAF, whereas ARAF displays the lowest intrinsic activity.(Chong & Guan, 2003; Cope et al., 2019) In line with this, RAF dimers exhibit a defined activity hierarchy: the BRAF-CRAF heterodimer is most active, followed by BRAF and CRAF homodimers, with ARAF-containing dimers generally least active. (Cope et al., 2018; Cope et al., 2020; Gunderwala et al., 2019; Yu et al., 2023) Within this framework, CRAF occupies a uniquely central position in RAS-MAPK signal transmission. Although activating mutations in CRAF are relatively rare, CRAF is indispensable for efficient MAPK signaling downstream of oncogenic RAS.(Cope et al., 2018; Montagut et al., 2008) CRAF forms catalytically potent heterodimers with BRAF, enhances BRAF’s affinity for HRAS, and contributes to adaptive resistance mechanisms, positioning it as a central amplifier of MAPK signaling.(Montagut et al., 2008; Villanueva et al., 2011; Villanueva et al., 2010) ARAF, though less characterized and catalytically weaker, has emerged as a context-dependent contributor to oncogenic signaling, consistent with reports of ARAF alterations in lung adenocarcinoma and pancreatic cancer. (Diamond et al., 2016; Imielinski et al., 2014)

RAS, a small GTPase, is positioned upstream in the MAPK signaling cascade and serves as a molecular switch that cycles between an inactive GDP-bound "off" state and an active GTP-bound "on" state.(Malumbres & Barbacid, 2003) HRAS, KRAS4a, KRAS4b, and NRAS maintain ∼80% sequence identity and possess identical effector-binding domains, allowing all four isoforms to interact with the same downstream effectors, yet they differ markedly in their membrane organization, trafficking, and oncogenic prevalence.(Furth et al., 1987; Leon et al., 1987) How the intrinsic differences in RAS isoforms translate into selective recruitment and activation of CRAF and ARAF, especially regarding the contributions of the CRD, remains insufficiently defined.

Targeting RAS and RAF proteins therapeutically has proven to be challenging, with issues related to the specificity, potency, and resistance mechanisms.(Davies et al., 2002; Durrant & Morrison, 2018) These challenges are largely due to gaps in our understanding of the complex molecular mechanisms governing the regulation of RAF by RAS and the intricacies of their interaction mechanisms within the MAPK signaling cascade. While cryo-EM structures have defined autoinhibited and activated RAF assemblies and illuminated the architecture of 14-3-3 stabilized complexes, these static snapshots have yielded divergent interpretations of whether RAS binding alone is sufficient to relieve autoinhibition, how the CRD is repositioned during activation, and how access to the kinase domain dimer interface is achieved. (Martinez Fiesco et al., 2022; Spencer-Smith & Morrison, 2024; Tran et al., 2005) Adding further complexity, RAS isoforms, despite sharing identical effector-binding surfaces, differ substantially in membrane organization and oncogenic context, yet how these differences are decoded by individual RAF isoforms remains poorly understood. Notably, NRAS remains the least mechanistically characterized of the major RAS isoforms despite its prominent role in melanoma and hematologic malignancies.

In this work, we address these fundamental gaps by defining how CRAF and ARAF N-terminal regulatory regions engage HRAS, KRAS4b, and NRAS. We elucidate the mechanistic contributions of the CRD to isoform-specific RAF recruitment and relief of autoinhibition. Through quantitative biophysical measurements, hydrogen-deuterium exchange mass spectrometry (HDX-MS), and analysis of structural interfaces, we uncover distinct regulatory principles that differentiate RAF isoforms and reveal CRD-dependent mechanisms that govern RAS-RAF complex formation and activation.

## Results

### CRAF^CRD^-dependent regulation drives isoform-selective RAS recognition

To delineate the contributions of individual domains within the CRAF N-terminal regulatory region to RAS recognition, we carried out a series of biophysical experiments to quantify the interactions between full-length HRAS, KRAS4b, and NRAS and CRAF constructs containing either the RBD alone or the combined RBD-CRD domains.

Purification of recombinant CRAF^RBD^, CRAF^CRD^, CRAF^RBD^ ^CRD^, and each RAS isoform (Figure 1A-B) enabled a controlled comparison of isoform-specific binding behaviors. RAS proteins were prepared in both their GDP-bound, inactive conformation and their GMPPNP-loaded, active state. CRAF constructs were incubated with GST-tagged RAS isoforms, and protein-protein interactions were assessed via GST pulldown assays. It is noteworthy that KRAS exists as two splice isoforms (KRAS4a and KRAS4b). KRAS4b is the predominant oncogenic form and the form most commonly used in biochemical studies, including in this study. As shown in Figure 1C-E, all three active RAS isoforms robustly interacted with both CRAF^RBD^ and CRAF^RBD^ ^CRD^ constructs, whereas GDP-loaded RAS displayed only weak or negligible binding. This clear discrimination between active and inactive RAS binding to the CRAF^RBD^ and CRAF^RBD^ ^CRD^ confirms nucleotide-state-dependent binding and validates the purified protein system. The CRAF^CRD^ construct had negligible binding to all three RAS isoforms in their GMPPNP bound form when compared to the GDP loaded RAS controls (Figure S1), suggesting CRD alone is not sufficient for binding. Therefore, this construct was excluded from subsequent studies.

**Figure 1:**
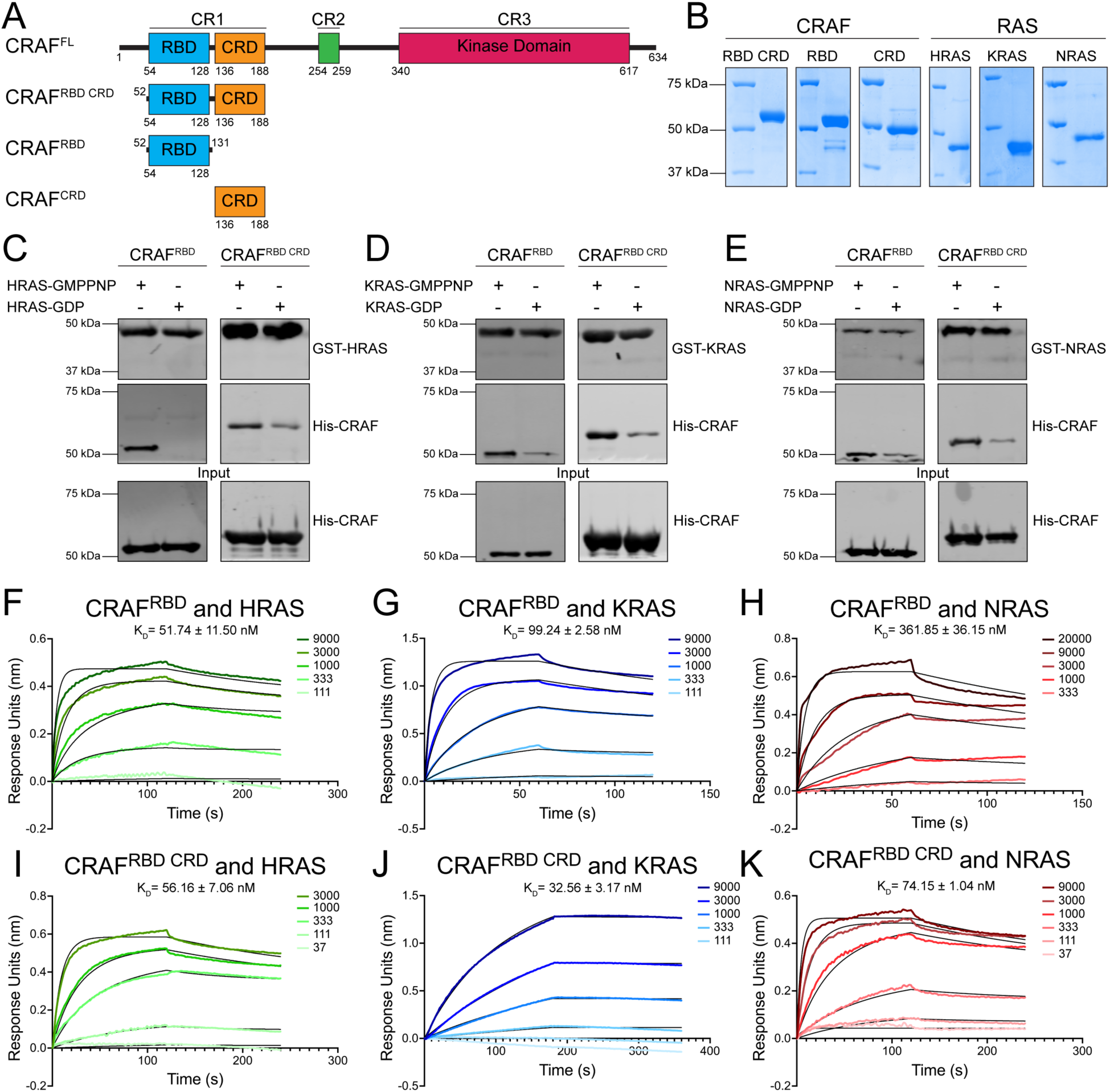
CRAF engages RAS isoforms with high nanomolar affinity through its N-terminal regulatory domains. A) Schematic representation of the regulatory architecture of full-length CRAF (CRAF^FL^) and the isolated N-terminal regulatory constructs used in this study. B) Coomassie-stained SDS-PAGE gels showing purified CRAF^RBD^ and CRAF^RBD^ ^CRD^ constructs along with H-/K-/NRAS. C-E) Pulldown analyses demonstrating preferential binding of CRAF^RBD^ and CRAF^RBD^ ^CRD^ to active HRAS (C), KRAS (D), or NRAS (E) in the GMPPNP-loaded state relative to GDP-bound RAS. Pulldown experiments were performed in two independent replicates with comparable results. (F–H) Quantitative BLI measurements showing nanomolar-affinity binding between CRAF^RBD^ and HRAS (F), KRAS (G), or NRAS (H). (I–K) BLI analysis of the CRAF^RBD^ ^CRD^ construct reveals enhanced nanomolar-affinity binding to HRAS (I), KRAS (J), and NRAS (K), highlighting a CRD-dependent stabilization of RAS engagement, particularly for NRAS. All BLI experiments were performed in duplicate. Association (*k_on_*) and dissociation (*k_off_*) rate constants were obtained by global fitting of the sensorgrams to a 1:1 binding model, and binding affinity (*K_D_*) was calculated as *k_off_*/*k_on_*. Fits exhibited high quality (R² > 0.9; χ² < 4).

We next assessed the differential RAS isoform selectivity of CRAF using the domain primarily responsible for RAS engagement, the RBD. The purified CRAF^RBD^ construct, we examined its binding kinetics across all three major RAS isoforms. Using biolayer interferometry (BLI), we observed that CRAF^RBD^ binds both HRAS and KRAS with low-nanomolar affinity (*K_D_* = 51.74 ± 11.50 nM and 99.24 ± 2.58 nM, respectively), establishing that the RBD is independently capable of forming a stable complex with H-and KRAS agreeing with previous isothermal titration calorimetry studies (Figure 1F-G).(Johnson et al., 2017) In contrast, NRAS binding was substantially weaker, with a *K_D_* of 361.85 ± 36.15 nM; nearly an order-of-magnitude decrease relative to HRAS and KRAS (Figure 1H). This marked reduction highlights an intrinsic limitation of the RBD in engaging NRAS and suggests that NRAS-CRAF recruitment may require additional stabilizing interactions beyond those provided by the RBD. Of note, while HRAS, KRAS, and NRAS all have similar association rates (*k_on_* ≈ 1.80×10^4^, 4.02×10^4^, and 1.15×10^4^ M^−1^ s^−1^, respectively); their dissociation rates slightly differ with HRAS having the slowest dissociation (*k_off_* ≈ 8.59×10^-4^ s^−1^) followed by KRAS (*k_off_* ≈ 4.01×10^-3^ s^−1^) then NRAS (*k_off_* ≈ 4.15×10^-3^ s^−1^).

A direct comparison of these measured affinities with previously reported values for CRAF RBD-H/KRAS interactions is provided in Supplementary Table S2, demonstrating overall agreement across studies while highlighting differences attributable to experimental approach and construct design.

Having established baseline contributions of the RBD, we next evaluated the effect of adding the CRD region using the purified CRAF^RBD^ ^CRD^ construct. Our findings revealed that CRAF^RBD^ ^CRD^ binds all three RAS isoforms with low nanomolar affinity, but with a hierarchy. KRAS exhibited the tightest interaction (*K_D_* = 32.56 ± 3.17 nM), followed by HRAS (*K_D_* = 56.16 ± 7.06 nM) and NRAS (*K_D_* = 74.15 ± 1.04 nM) (Figure 1I-K). These differences, although modest in magnitude, establish an intrinsic gradient of isoform preference within the context of the full N-terminal regulatory domain. Inclusion of the CRD enables KRAS to engage CRAF most efficiently despite having the slowest association rate (*k_on_* ≈ 1.04×10^3^ M^−1^ s^−1^), it also has an extremely slow dissociation rate (*k_off_* ≈ 3.15×10^-5^ s^−1^) indicating that KRAS forms a more stable complex with CRAF, when the CRD is present, relative to HRAS and NRAS. Interestingly, the contribution of the CRD to HRAS binding appeared minimal. The CRAF^RBD^ ^CRD^-HRAS interaction remained in the same affinity range as the RBD alone construct, suggesting that HRAS-CRAF engagement is driven predominantly by the RBD. (Figure 1F and I). In contrast, NRAS binding was highly sensitive to the presence of the CRD. CRAF^RBD^ ^CRD^ bound NRAS with markedly increased affinity, representing nearly a five-fold enhancement compared with the RBD construct (Figure 1H and K). Thus, among the three isoforms, NRAS uniquely depends on CRD-mediated contacts for productive engagement.

### ARAF interprets RAS isoforms through differential engagement of its N-terminal regulatory architecture

We next sought to characterize the molecular determinants governing ARAF recruitment with distinct RAS isoforms. Although ARAF remains the least mechanistically understood RAF family member, emerging evidence suggests that it exhibits regulatory properties distinct from those of BRAF and CRAF. (Tran et al., 2005) To investigate this, we evaluated the binding properties of isolated ARAF N-terminal domains, (Figure 2A) using a combination of pulldown assays and BLI. Coomassie-stained SDS-PAGE gels (Figure 2B) confirm high-purity isolation of ARAF^RBD^ and ARAF^RBD^ ^CRD^ constructs, providing suitable material for quantitative binding studies.

**Figure 2:**
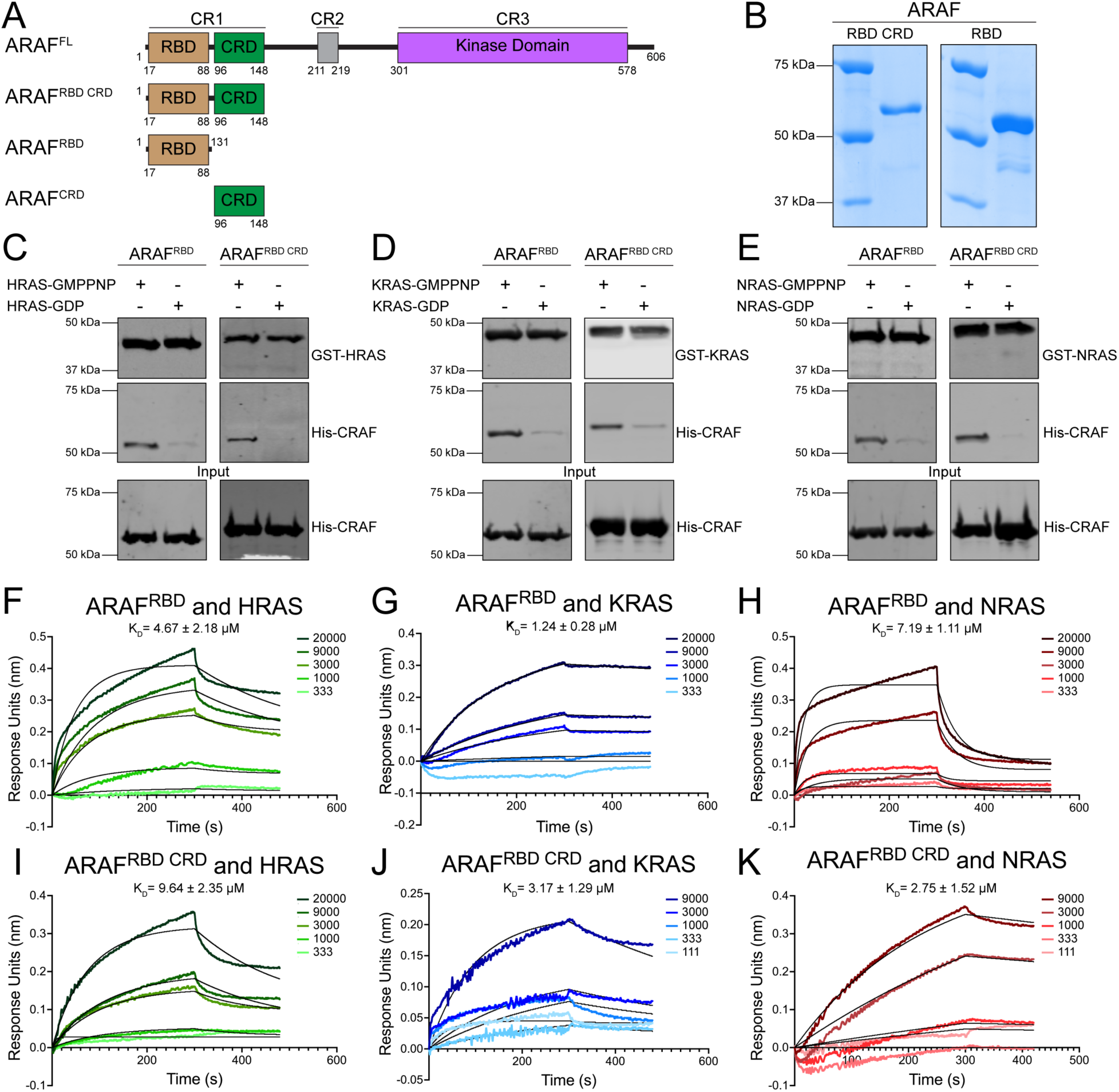
ARAF engages RAS with micromolar affinity through its N-terminal regulatory domains. A) Depiction of ARAF regulatory domains within full-length ARAF (ARAF^FL^) and N-terminal isolated regulatory constructs of ARAF. B) Coomassie stained gels of purified ARAF^RBD^ and ARAF^RBD^ ^CRD^, C-E) Pulldown analysis demonstrating preferential binding of ARAF^RBD^ and ARAF^RBD^ ^CRD^ to active HRAS- (C), KRAS- (D), or NRAS- (E) GMPPNP relative to GDP-bound RAS. Pulldown experiments were performed in two independent replicates with comparable results. F-H) Quantitative BLI analysis of binding between ARAF^RBD^ and HRAS (F), KRAS (G), or NRAS. I-K) BLI analysis of the ARAF^RBD^ ^CRD^ construct reveals HRAS µM) (I), KRAS (J), or NRAS (K) binding in the micromolar range. All BLI experiments were performed in duplicate. Association (*k_on_*) and dissociation (*k_off_*) rate constants were obtained by global fitting of the sensorgrams to a 1:1 binding model, and binding affinity (*K_D_*) was calculated as *k_off_*/*k_on_*. Fits exhibited high quality (R² > 0.9; χ² < 4).

Similar to our previous analysis with CRAF, pulldown assays were performed using GST-tagged HRAS, KRAS, or NRAS loaded with either GMPPNP (active) or GDP (inactive). Across all three isoforms, active RAS species displayed robust interaction with both ARAF constructs, whereas GDP-bound RAS showed minimal or undetectable binding (Figure 2C-E). These results validate that ARAF engagement is strongly dependent on the active, GTP-like conformation of RAS.

We first examined the binding properties of ARAF^RBD^ alone, enabling the isolated contribution of the RBD to be assessed (Figure 2F-H). The ARAF^RBD^ construct binds HRAS with a *K_D_* of 4.67 ± 2.18 μM, indicating relatively weak affinity. This is consistent with the low intrinsic catalytic efficiency of ARAF and suggests limited stabilizing contacts within the ARAF^RBD^-HRAS interface (Figure 2F). The affinity for KRAS (*K_D_* = 1.24 ± 0.28 μM) is stronger than for HRAS but still in the micromolar range (Figure 2G). Importantly, even KRAS, typically the strongest RAS isoform for RAF engagement, binds ARAF^RBD^ with substantially weaker affinity compared to CRAF^RBD^, highlighting inherent isoform-specific differences in RBD engagement between RAF isoforms (Figure 1G). Binding to NRAS is the weakest among the three (*K_D_* = 7.19 ± 1.11 μM).

This suggests NRAS is the poorest intrinsic recruiter of ARAF when only the RBD is present, mirroring NRAS’s consistently weaker interactions, as seen with CRAF (Figure 1H and 2H). Collectively, these results reveal that the ARAF^RBD^ exhibits overall low affinity toward all RAS isoforms, with a clear isoform-selective ranking: KRAS > HRAS > NRAS.

We next assessed the influence of the CRD by analyzing binding of the ARAF^RBD^ ^CRD^ construct. Surprisingly, inclusion of the CRD weakens affinity toward HRAS (*K_D_* = 9.64 ± 2.35 μM), nearly doubling the *K_D_* value relative to the RBD alone. This indicates that the CRD does not contribute stabilizing contacts in the ARAF-HRAS interface and may even induce an orientation that is suboptimal for HRAS engagement (Figure 2I). KRAS binding also weakens upon CRD inclusion (*K_D_* = 3.17 ± 1.29 μM vs. 1.24 ± 0.28 μM for RBD alone) (Figure 2J). This contrasts sharply with CRAF, where the CRD nearly doubled KRAS affinity (Figure 1J). Thus, while the CRD in CRAF cooperates with the RBD to stabilize engagement, the CRD in ARAF exerts the opposite effect, slightly destabilizing HRAS and KRAS binding. In contrast to the pattern with HRAS and KRAS, inclusion of the CRD in ARAF results in a modest but reproducible improvement in NRAS binding affinity (*K_D_* 2.75 ± 1.52 μM vs. 7.19 ± 1.11 μM for RBD alone), corresponding to an approximately 2.6-fold enhancement (Figure 2K). While this effect is less pronounced than the CRD-dependent stabilization observed for CRAF, it highlights a distinct and isoform-specific contribution of the ARAF CRD to NRAS engagement. This isoform-dependent behavior reveals that the ARAF CRD performs divergent roles depending on the RAS isoform: destabilizing for HRAS and KRAS but stabilizing for NRAS.

According to prior structural studies, the majority of previously identified KRAS-contacting residues are conserved between CRAF and ARAF, with 11 of 12 residues conserved in the RBD and 10 of 13 residues conserved in the CRD (Figure S2).(Tran et al., 2021) Notably, the KRAS interacting residues within RBD occur in conserved clusters, whereas, several non-conserved residues cluster around the conserved CRD-KRAS interaction surface, suggesting that differences in the local chemical environment, CRD positioning, or RBD-CRD coupling may contribute to the divergent CRD-dependent effects observed for CRAF and ARAF.

### CRAF CRD Provides Critical Stabilization for NRAS Binding

Our quantitative binding measurements indicated that the CRAF CRD plays a disproportionate role in stabilizing NRAS engagement, yielding an approximately five-fold increase in affinity for the CRAF^RBD^ ^CRD^ construct relative to the RBD alone. To define the structural basis of this effect, we employed HDX-MS to interrogate domain-specific contributions to NRAS recognition.

We first examined the isolated CRAF^RBD^ to determine whether the RBD alone is sufficient to stabilize NRAS binding. Comparison of CRAF^RBD^ in the apo state versus in the presence of NRAS revealed no appreciable reduction in deuterium uptake across the RBD region (Figure 3A), indicating that NRAS engagement does not measurably stabilize the RBD in isolation. Notably, although BLI measurements detect a sub-micromolar interaction between CRAF^RBD^ and NRAS, HDX-MS reports on ensemble-averaged conformational protection and is inherently less sensitive to transient, rapidly dissociating complexes. Given the relatively fast dissociation kinetics of the CRAF^RBD^-NRAS interaction, any protection arising from this interaction is likely averaged out during the labeling period and falls below the threshold for confident detection.

**Figure 3:**
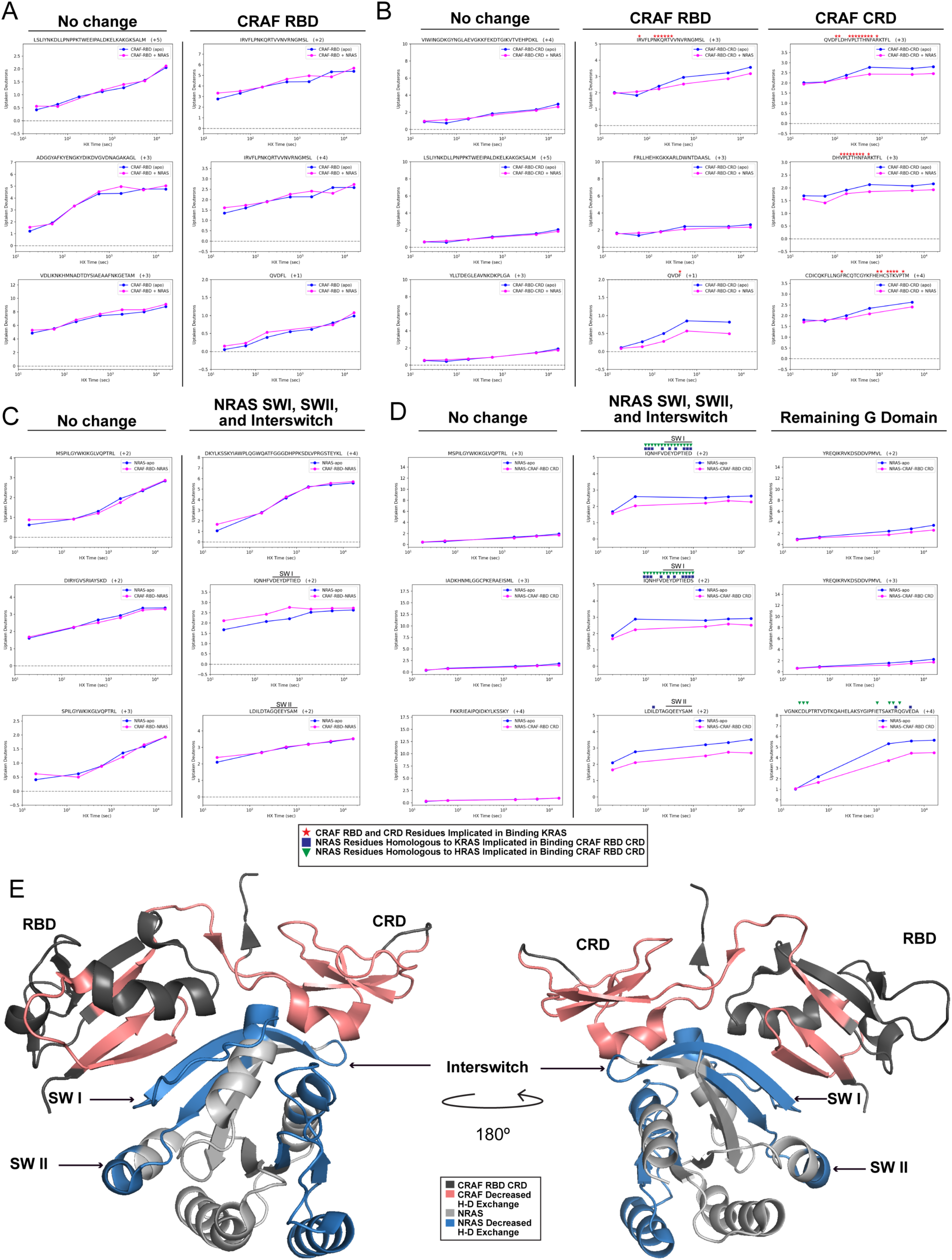
The CRAF cysteine-rich domain stabilizes NRAS engagement through a cooperative interface. A) Representative HDX-MS plots of CRAF^RBD^ peptides showing no appreciable protection from deuterium uptake in CRAF^RBD^ in the absence (blue) and presence (pink) of NRAS. B) In contrast, the CRAF^RBD^ ^CRD^ construct exhibits slowed deuterium uptake across the RBD and CRD regions in the presence (pink) of NRAS compared with the absence (blue) of NRAS. CRAF residues previously implicated in KRAS interactions are indicated with red stars. C) Representative HDX-MS plots of NRAS peptides indicates no significant changes in deuterium incorporation within the switch I, switch II, or interswitch regions in the absence (blue) versus presence (pink) of CRAF^RBD^. D) Representative HDX-MS plots of NRAS peptides reveals corresponding decreases in deuterium uptake within switch I, switch II, and interswitch regions when CRAF^RBD^ ^CRD^ is present (pink) compared to when CRAF^RBD^ ^CRD^ is absent (blue) consistent with formation of a stabilized, cooperative interface. NRAS residues homologous to KRAS (blue squares) and HRAS (green triangle) residues previously implicated in binding CRAF^RBD^ ^CRD^ have been indicated. E) CRAF^RBD^ ^CRD^ (PDB: 6XHB) in complex with NRAS (PDB: 5UHV) regions with slowed deuterium uptake are shown in pink for CRAF and blue for NRAS. Key interacting regions within NRAS such as, SW I, SW II, and interswitch regions have been indicated in the structure.

Consistent with this observation, reciprocal HDX-MS analysis of NRAS showed no significant changes in deuterium uptake within the switch I, switch II, or interswitch regions when NRAS was incubated with CRAF^RBD^ alone (Figure 3C). Together, these data indicate that the canonical RBD-switch interaction is insufficient to generate a stable CRAF-NRAS complex in the absence of the CRD.

In contrast, inclusion of the CRD produced pronounced stabilization on both binding partners. Comparison of CRAF^RBD^ ^CRD^ in the apo state versus in the presence of NRAS revealed substantial protection from deuterium uptake across regions spanning both the RBD and CRD (Figure 3B). These protected segments map to surfaces previously implicated in KRAS engagement.(Tran et al., 2021) Structural mapping of these residues onto the CRAF^CRD^ highlights a surface that overlaps with the experimentally defined KRAS-CRD interaction interface and includes residues such as H139, R143, F163, E174, S177, and T178, which are known to participate in hydrogen-bonding interactions with KRAS.(Tran et al., 2021) Importantly, prior mutagenesis studies targeting residues within this region, particularly R143 and F163, have demonstrated that perturbations of this CRD surface can alter CRAF activation and RAS-dependent signaling, underscoring its functional relevance.(Cutler & Morrison, 1997; Winkler et al., 1998)

Reciprocal HDX-MS analysis further supported this model. Upon binding to CRAF^RBD^ _CRD_, NRAS exhibited marked reductions in deuterium uptake within the switch I, switch II, and interswitch regions, as well as at the C-terminal portion of the G-domain proximal to the hypervariable region (Helix 4) (Figure 3D and S5). Notably, peptides corresponding to the interswitch region (residues 40-48) exhibit clear protection in the presence of CRAF^RBD^ ^CRD^, consistent with stabilization of this region upon complex formation as shown in previous CRAF^RBD^ ^CRD^-HRAS work (Figure S6). (Cookis & Mattos, 2021) Helix 4 (α4), while distant from the canonical switch I/II effector interface, resides within the allosteric lobe of the G-domain. Prior NMR and MD simulation studies with HRAS have shown this allosteric region to undergo long-range conformational rearrangements upon effector (CRAF) binding at the switch regions.(Buhrman et al., 2010; Cookis & Mattos, 2021; Fetics et al., 2015) Peptide coverages for all apo state proteins were determined to be >95% (Figure S3-S5). Structural mapping of these protected regions onto the NRAS crystal structure reveals a broadened interaction surface that extends beyond the canonical switch regions, providing a mechanistic explanation for the enhanced affinity observed in the presence of the CRD.

### KRAS Binding Weakens the Interaction Between CRAF N-terminal Regulatory Domains and the Kinase Domain

Structural comparison of the CRAF^CRD^ in its autoinhibited and active states reveals a clash within its binding interfaces and underscores the CRD as a key regulator of RAF activation. In the autoinhibited conformation (Figure 4A and B), CRD residues R143, L147, L149, F151, K157, F158, L160, and N161 engage the 14-3-3 dimer, while K148, L149, F150, D153, Q156, T167, G169, and K171 interact with the CRAF^KD^ forming a composite interface that stabilizes the closed, monomeric assembly and limits access to the dimer interface.(Park et al., 2019) In the active conformation (Figure 4C), residues involved in membrane association and KRAS engagement (L147, K148, L149, A150, K157, and F158) substantially overlap with those that contact 14-3-3 in the autoinhibited state. The structural overlay (Figure 4D) demonstrates that these residues cannot accommodate both interaction modes simultaneously, indicating that RAS binding directly competes with autoinhibitory CRD interactions.

**Figure 4:**
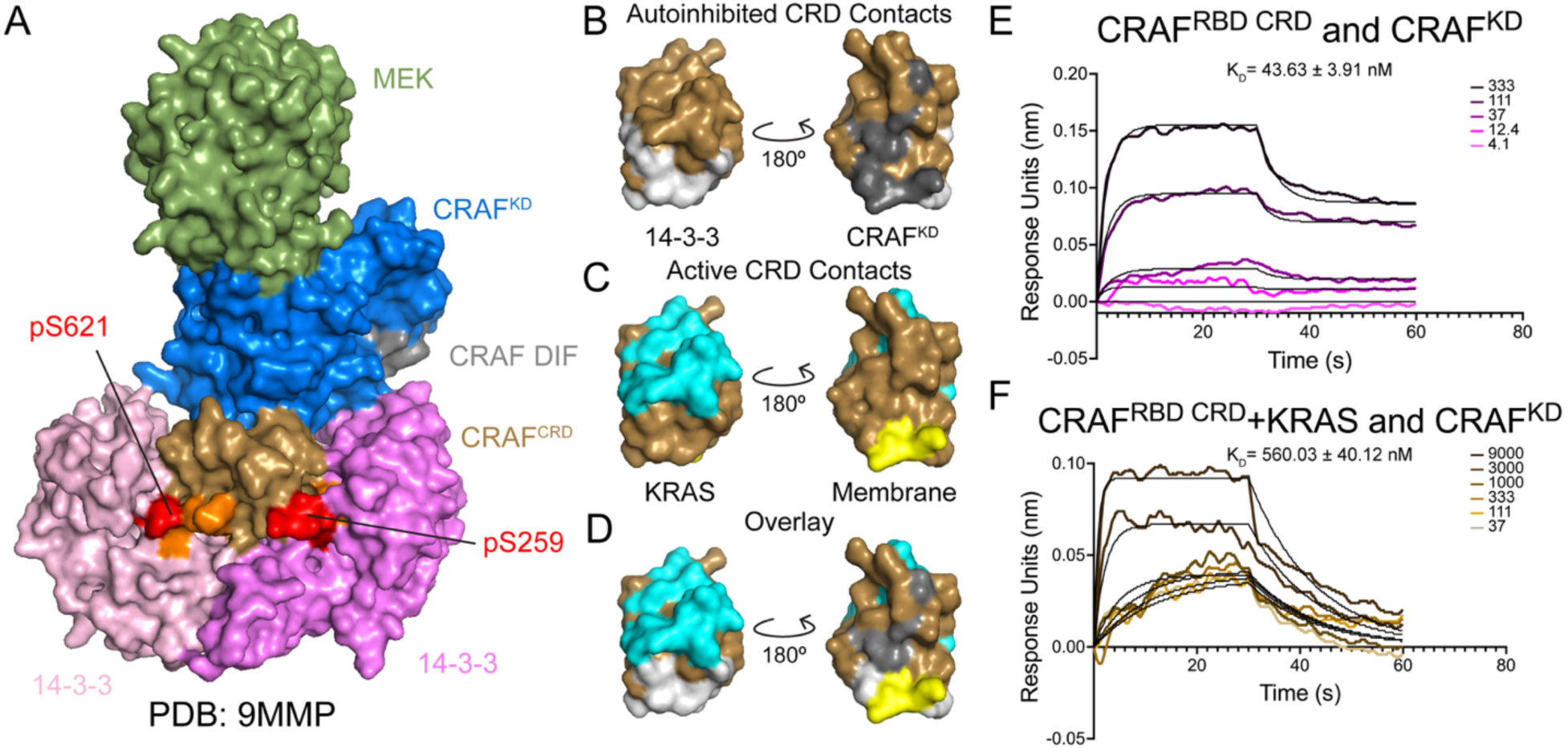
KRAS binding directly impacts CRAF N- and C-terminal autoinhibitory interactions. A) Structure of autoinhibited monomeric CRAF bound to MEK and 14-3-3 protomers. B) Isolated CRAF^CRD^ with autoinhibitory 14-3-3 contacts highlighted in white and CRAF^KD^ contacts highlighted in grey. C) Isolated CRAF^CRD^ with activating KRAS contacts highlighted in cyan and membrane contacts highlighted in yellow. D) Isolated CRAF^CRD^ with autoinhibitory 14-3-3 and CRAF^KD^ contacts overlayed with activating KRAS and membrane contacts. E) Quantitative BLI analysis of binding between CRAF^RBD^ ^CRD^ and CRAF^KD^. F) Quantitative BLI analysis of binding between CRAF^RBD^ _CRD_ and CRAF^KD^ in the presence of KRAS. All BLI experiments were performed in duplicate. Association (*k_on_*) and dissociation (*k_off_*) rate constants were obtained by global fitting of the sensorgrams to a 1:1 binding model, and binding affinity (*K_D_*) was calculated as *k_off_*/*k_on_*. Fits exhibited high quality (R² > 0.9; χ² < 4).

We next investigated whether RAS engagement with the CRAF N-terminal regulatory region (CRAF^RBD^ ^CRD^) directly perturbs autoinhibitory interactions between the N- and C-terminal domains of CRAF. We began by studying the binding between CRAF^RBD^ ^CRD^ and CRAF^KD^ using BLI. Our findings revealed a remarkably tight intramolecular interaction (*K_D_* = 43.63 ± 3.91nM), consistent with a highly stable autoinhibited configuration in which the N-terminal domains strongly associate with the kinase domain (Figure 4E). Next, we pre-incubated CRAF^RBD^ ^CRD^ with KRAS in a 1:1 molar ratio for one hour and repeated the binding analysis with CRAF^KD^. The presence of KRAS resulted in an over 10-fold reduction in affinity for CRAF^KD^ (*K_D_* = 560.03 ± 40.12 nM), demonstrating that KRAS binding substantially weakens these autoinhibitory contacts. Taken together, these results demonstrate that KRAS binding to the isolated CRAF^RBD^ _CRD_ construct reduces its affinity for the CRAF kinase domain. Although this simplified reconstituted system does not recapitulate the full autoinhibited RAF complex, which includes full-length RAF, 14-3-3, membrane context, and native post-translational modifications, the data suggest that RAS engagement is incompatible with simultaneous stabilization of this CRAF^RBD^ ^CRD^-CRAF^KD^ interaction.

### KRAS inhibitors substantially decrease binding between CRAF and K/NRAS

To assess how emerging KRAS-targeted inhibitors influence early RAS-RAF interactions, we examined their effects on binding between the CRAF N-terminal regulatory region, CRAF^RBD^ ^CRD^ and individual RAS isoforms using BLI. All inhibitor BLI experiments of KRAS were performed using full-length KRAS4b loaded with GMPPNP, allowing inhibitor effects to be evaluated in the context of an active, GTP-like RAS substrate. We first evaluated BI 2865, a noncovalent pan-KRAS inhibitor that binds an allosteric pocket formed by the switch I and switch II regions of KRAS.(Kim et al., 2023) By stabilizing KRAS in an inactive conformation, BI 2865 acts by disrupting effector interactions.(Kim et al., 2023; Tedeschi et al., 2025) We sought to investigate how this inhibitor affects KRAS-CRAF binding (Figure 5A). GMPPNP-loaded full-length KRAS4b was pre-incubated with BI 2865 for 1 hr at 4 °C prior to BLI analysis, after which binding to CRAF^RBD^ ^CRD^ was measured under conditions identical to those used for prior CRAF and ARAF experiments. The presence of BI 2865 resulted in an approximately 20-fold reduction in binding affinity between CRAF^RBD^ ^CRD^ and KRAS, indicating substantial disruption of the KRAS-RAF interface.

**Figure 5:**
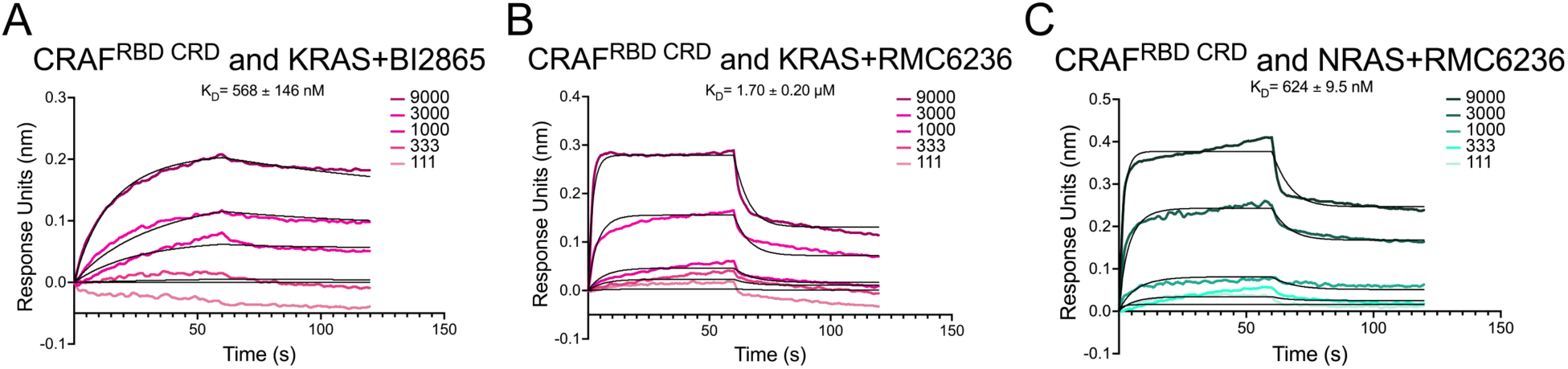
KRAS inhibitors directly perturb the RAS-CRAF interface. Quantitative BLI analysis of binding between CRAF^RBD^ ^CRD^ and KRAS in the presence of A) BI 2865, B) RMC 6236, and C) Quantitative BLI analysis of binding between CRAF^RBD^ ^CRD^ and NRAS in the presence of RMC 6236. All BLI experiments were performed in duplicate. Association (*k_on_*) and dissociation (*k_off_*) rate constants were obtained by global fitting of the sensorgrams to a 1:1 binding model, and binding affinity (*K_D_*) was calculated as *k_off_*/*k_on_*. Fits exhibited high quality (R² > 0.9; χ² < 4).

We next examined RMC 6236, a noncovalent, pan-RAS(ON) inhibitor that selectively engages the active, GTP-bound state of multiple RAS isoforms by binding a conserved pocket adjacent to the switch II region.(Jiang et al., 2024) This interaction sterically and allosterically blocks effector binding, thereby suppressing downstream signaling across diverse RAS mutants (Cregg et al., 2025; Jiang et al., 2024). Pre-incubation of KRAS with RMC 6236 led to a more pronounced effect, producing an approximately 56-fold decrease in affinity for CRAF^RBD^ ^CRD^ (Figure 5B). Given the broader specificity of RMC 6236, we additionally assessed its impact on the CRAF-NRAS interaction. Consistent with its pan-RAS activity, RMC 6236 reduced binding between CRAF^RBD^ ^CRD^ and NRAS by approximately 8-fold (Figure 5C).

In the case of CRAF-KRAS, BI 2865 increases the association rate approximately 10-fold (*k_on_* ≈ 1.1 × 10⁴ M⁻¹ s⁻¹) but also elevates the dissociation rate by nearly two orders of magnitude (*k_off_* ≈ 3.1 × 10⁻³ s⁻¹), substantially shortening complex lifetime. RMC 6236 produces an even more pronounced kinetic shift, accelerating association by roughly 50-fold (*k_on_* ≈ 5.3 × 10⁴ M⁻¹ s⁻¹) while increasing dissociation by more than three orders of magnitude (*k_off_* ≈ 7.9 × 10⁻² s⁻¹). A similar kinetic pattern is observed for CRAF-NRAS, where RMC 6236 substantially accelerates *k_off_* (≈4.2×10^−2^ versus 1.8×10^−3^ s^−1^) relative to the uninhibited state, with KRAS displaying greater overall sensitivity. Together, these results demonstrate that clinically relevant KRAS and pan-RAS(on) inhibitors substantially weaken CRAF-RAS interactions, highlighting their capacity to perturb early steps in RAF recruitment and activation.

### Small-Molecule Targeting of RAS Perturbs CRAF-RAS Interaction in a Cellular Context

To independently validate the effects of RAS inhibitors on CRAF-RAS interactions in a cellular context, we developed a NanoBiT-based complementation assay to monitor RAS-CRAF association in live cells (Figure 6A). In this system, CRAF and RAS were fused to the small (SmBiT) and large (LgBiT) subunits of the NanoBiT luciferase, respectively, such that reconstitution of enzymatic activity upon protein-protein interaction produced a luminescence signal proportional to complex formation. RAS-LgBiT and CRAF-SmBiT constructs were coexpressed in HEK293 cells, and luminescence was measured following treatment with a panel of clinically relevant KRAS inhibitors. These included the covalent KRAS^G12C^ inhibitors Sotorasib and Adagrasib, which irreversibly modify the mutant cysteine within the switch II pocket to stabilize KRAS in its inactive GDP-bound state, as well as BI 2865 and RMC 6236.

**Figure 6:**
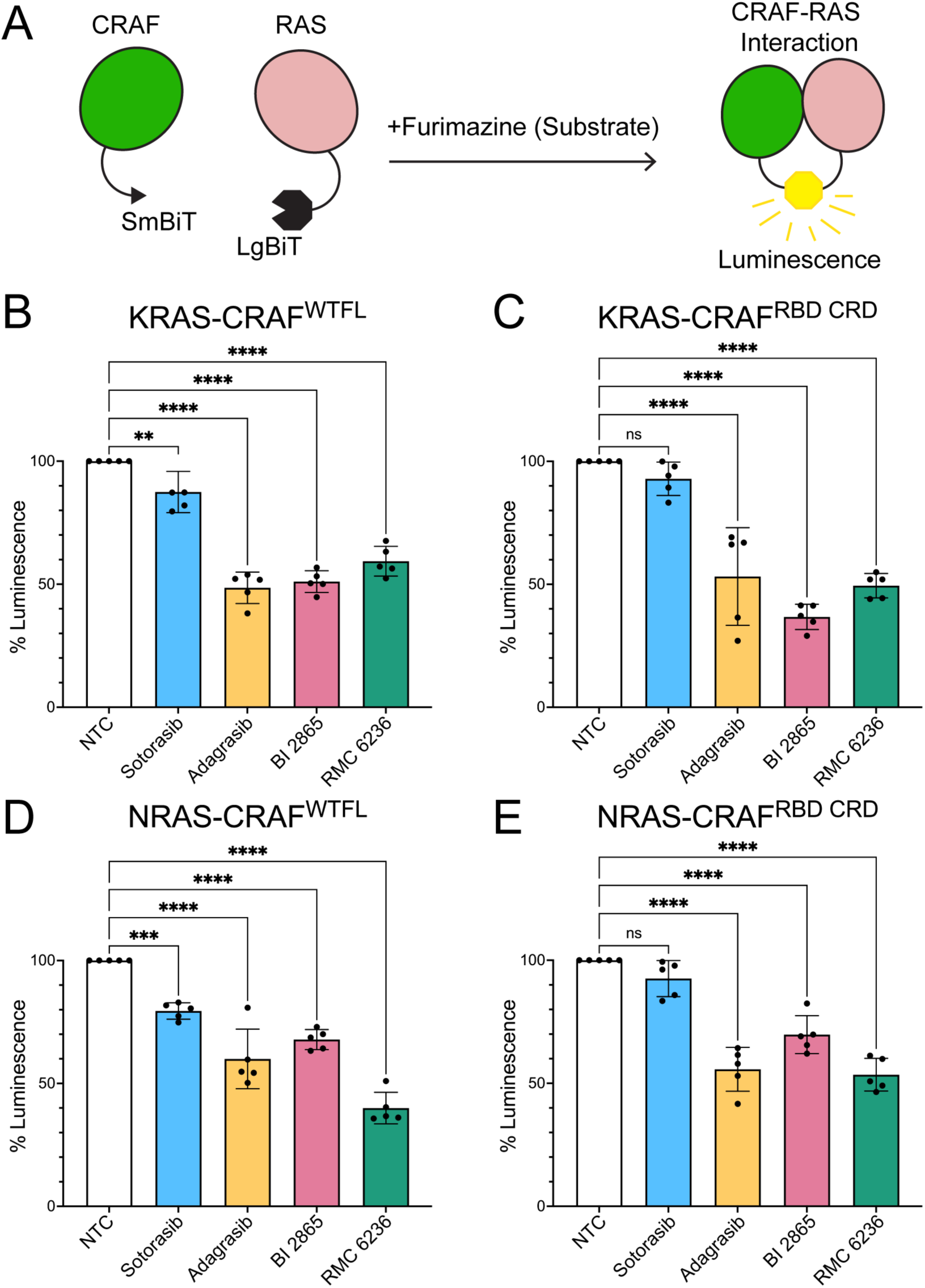
KRAS and pan-RAS inhibitors disrupt CRAF–RAS interactions in live cells. A) Schematic of the NanoBiT complementation assay used to monitor CRAF-RAS interactions in live cells. HEK293 cells expressing respective NanoBiT constructs were treated with 10µM of indicated RAS inhibitors for 4 hrs. B) NanoBiT analysis of CRAF^WTFL^ interaction with KRAS in HEK293 cells following treatment with clinically relevant KRAS and pan-RAS inhibitors, Sotorasib, Adagrasib, BI 2865, and RMC 6236. B) NanoBiT analysis of the CRAF^RBD^ ^CRD^ N-terminal regulatory region interacting with KRAS, in the absence and presence of Sotorasib, Adagrasib, BI 2865, and RMC 6236. D-E) Extension of the NanoBiT assay to CRAF-NRAS interaction in response to RAF inhibitor treatment. All NanoBiT assays were normalized to no treatment control across five biological replicates. Statistical significance was determined through paired t test where indicated, followed by Tukey’s honest significant different post-hoc test with corresponding P-values (*P<0.05, **P<0.01, ***P<0.001). All graph bars represent the mean ± SD with individual data points per biological replicate with corresponding p-values.

Using full-length wild-type CRAF (CRAF^WTFL^), we observed a pronounced reduction in luminescence upon treatment with Adagrasib, BI 2865, and RMC 6236, corresponding to an approximately 50% decrease in signal relative to untreated controls, indicating substantial disruption of the CRAF^WTFL^ -KRAS interaction in cells (Figure 6B). In contrast, Sotorasib produced only a modest decrease in signal of about 10% under identical conditions. We next examined whether similar effects were observed when the interaction was restricted to the N-terminal regulatory region of CRAF (Figure 6C).

Consistent with the full-length protein, the CRAF^RBD^ ^CRD^ construct exhibited marked sensitivity to Adagrasib and RMC 6236 (each producing ∼50% reductions in luminescence), while BI 2865 induced an even greater disruption (∼65% reduction). As observed for full-length CRAF, Sotorasib treatment did not significantly alter luminescence (∼10% reduction) relative to untreated controls. Expression levels of these proteins were confirmed to ensure that the reduction in luminescence is the result of disruption rather than degradation (Figure S7).

We further extended this analysis to NRAS to assess whether inhibitor effects were isoform dependent (Figure 6D-E). For the CRAF^WTFL^-NRAS interaction, Adagrasib and BI 2865 reduced luminescence by approximately 40%, whereas RMC 6236 produced a more pronounced effect (∼60% reduction). A similar trend was observed for the CRAF^RBD^ ^CRD^-NRAS interaction, with Adagrasib and RMC 6236 reducing luminescence by ∼45%, while BI 2865 elicited a more modest decrease (∼30%). For both the CRAF^WTFL^-NRAS and CRAF^RBD^ ^CRD^-NRAS, Sotorasib elicits only a modest reduction of ∼20% or negligible reduction in luminescence, respectively. Together, these cellular data corroborate the biophysical findings and demonstrate that these KRAS inhibitors can attenuate CRAF-RAS complex formation in cells. Moreover, the differential responses observed across inhibitors and RAS isoforms underscore that small-molecule engagement of RAS can variably perturb early RAF recruitment events, with implications for isoform-specific regulation of MAPK signaling.

## Discussion

By integrating quantitative biophysical measurements, HDX-MS, structural analysis, and live-cell protein-protein interaction assays, this study defines the mechanistic basis for isoform-specific RAS-RAF interactions. Together with our previous analysis of BRAF regulatory domains (Trebino et al., 2023), these findings establish a comparative framework for understanding how all three RAF isoforms decode RAS identity during early activation steps.

A major finding is that CRAF and ARAF interpret RAS isoform identity through distinct binding preferences. CRAF binds HRAS, KRAS, and NRAS with uniformly high nanomolar affinity, whereas ARAF binds all three isoforms substantially more weakly, in the low micromolar range. Despite their markedly different overall affinities, both RAF kinases display NRAS-specific behavior: in CRAF, NRAS binding is disproportionately dependent on CRD engagement, whereas in ARAF the CRD selectively stabilizes NRAS but destabilizes interactions with HRAS and KRAS. These results demonstrate that RAS isoforms are not functionally interchangeable and suggest that each RAF isoform has evolved distinct mechanisms for interpreting RAS identity.

Our findings are best understood in the context of prior BRAF studies. Previous work showed that although the BRAF RBD mediates high-affinity RAS binding, the BRAF-specific region (BSR) and CRD impose additional regulatory constraints that slow association kinetics and confer preference for KRAS over HRAS (Trebino et al., 2023). Structural studies of autoinhibited BRAF further revealed that the RBD is positioned for RAS engagement, whereas the CRD participates in intramolecular contacts that stabilize the inactive conformation (Martinez Fiesco et al., 2022). Together with the present data, these observations support a conserved model in which all RAF isoforms engage RAS through a shared RBD interface, while isoform-specific N-terminal regulatory architectures tune selectivity and activation.

HDX-MS provides structural insight into the basis of CRAF selectivity. NRAS binding induces cooperative protection across both the CRAF RBD and CRD, consistent with formation of an extended interface not observed for the isolated RBD. Reciprocal HDX-MS of NRAS reveals protection in the expected switch I, switch II, and interswitch regions. Additionally, increased deuterium uptake is also seen in helix 4 (α 4), distal to the canonical effector-binding site. This pattern suggests allosteric redistribution of conformational dynamics within NRAS, in which stabilization of switch regions is coupled to increased flexibility in the α 4/ α 5 lobe. Similar long-range communication has been reported in HRAS-CRAF complexes and supports a model in which effector binding remodels the global RAS conformational ensemble rather than only the local interface (Buhrman et al., 2010; Cookis & Mattos, 2021; Fetics et al., 2015; Tran et al., 2021). Loss of these protection signatures upon CRD deletion is consistent with the reduced affinity measured by BLI and indicates that stable NRAS-CRAF engagement requires CRD-dependent reinforcement of the RBD-switch interface.

The CRD-dependent requirement for stable NRAS-CRAF engagement may have particular relevance in NRAS-driven disease. NRAS mutations are enriched in melanoma and hematologic malignancies, tumor types frequently reported to depend on CRAF signaling despite the higher intrinsic catalytic activity of BRAF.(Akbani et al., 2015; Muñoz-Couselo et al., 2017) In contrast to KRAS-driven systems, which often rely on strong catalytic output supported by BRAF-containing dimers, NRAS-driven tumors may be more dependent on mechanisms that stabilize CRAF recruitment and activation. Our data suggest that CRD-mediated stabilization may represent one such regulatory dependency.

Our data also provide biochemical evidence that RAS engagement destabilizes intramolecular autoinhibitory contacts within CRAF. Structural comparisons of inactive and active RAF states indicate that CRD surfaces required for RAS and membrane binding overlap with interfaces that contact 14-3-3 and the kinase domain in autoinhibited complexes. (Martinez Fiesco et al., 2022; Park et al., 2019; Park et al., 2023) Consistent with this model, KRAS binding reduced affinity between the isolated CRAF RBD-CRD regulatory module and kinase domain by more than 10-fold, indicating that RAS engagement weakens N- to C-terminal interactions. These findings complement prior studies showing that RAS can bind autoinhibited RAF through the RBD while the CRD remains sequestered, and that membrane lipids further promote activation. (Park et al., 2023; Ritt et al., 2025) Together, our results support a multistep model in which RAS binding destabilizes autoinhibition but full activation additionally requires membrane recruitment, CRD repositioning, 14-3-3 rearrangement, and native post-translational regulation.

Our inhibitor studies demonstrate that emerging noncovalent and covalent RAS inhibitors profoundly destabilize the primary structural interface between RAS and the CRAF regulatory region, thereby halting early MAPK signalosome assembly. Strikingly, biophysical kinetic analyses revealed that noncovalent inhibitors (such as BI 2865 and RMC 6236) do not merely prevent protein association; rather, they paradoxically accelerate the association rate while massively increasing the dissociation rate by up to three orders of magnitude. This forces a highly unstable "hit-and-run" interaction whose drastically abbreviated lifetime precludes the prolonged membrane dwell time required for productive RAF activation. We confirmed this kinetic uncoupling in live cells using a NanoBiT complementation assay, which showed that both noncovalent inhibitors and the covalent inhibitor Adagrasib—but notably not Sotorasib—robustly dismantle CRAF-RAS complexes. Together, these findings establish a precise kinetic mechanism for how small-molecule engagement of the RAS switch pockets physically uncouples effector recruitment, while highlighting critical pharmacological differences among clinical inhibitors that may dictate therapeutic efficacy.

Because RAF activation is intrinsically membrane-coupled, these solution-phase measurements should be interpreted within the broader context of membrane-dependent signaling, where the CRD coordinates interactions with both RAS and membrane lipids. (Cookis & Mattos, 2021; Jimenez Salinas et al., 2026; Tran et al., 2021) By isolating protein–protein contributions in the absence of membrane, our system resolves isoform-specific CRD effects that are otherwise difficult to separate from membrane contributions. Future studies will be required to determine how these solution-phase differences are modulated in membrane-reconstituted systems.

## Methods and Materials

### Plasmids

GST-HRAS and His/MBP-KRAS were purchased from Addgene (#55653 and #159546). GST-KRAS was created with standard Gibson Assembly (NEB) procedures with pGEX2T as the vector and the following primers: 5’ATCTGGTTCCGCGTGGATCCACTGAATATAAACTTGTGGTAG3’(GST-KRAS_For), 5’CAGTCAGTCACGATGAATTCTTACATAATTACACACTTTGTCTTTG 3’ (GST-KRAS_Rev), 5’ GAATTCATCGTGACTGACTGACG 3’ (GST-vector_For), 5’ GGATCCACGCGGAACCAG 3’ (GST-vector_Rev). His/Halotag-CRAF RBD, His/MBP-CRAF RBD CRD, and His/MBP-CRAF CRD were purchased from Addgene (#159696, #159697, and #159698). 6X-HIS-BRAF-WT/FLAG or V5 was prepared as previously described, and MBP-CRAF-FLAG was created using common cloning procedures with pcDNATM 4/TO (Invitrogen) as the vector. CDC37-myc-FLAG was purchased from Origene (RC201002).

### Protein Expression

All protein construct plasmids were transformed into BL21 codon + E. coli. and grown to an OD600 0.6-0.8 in LB broth (CRAF RBD CRD and CRAF CRD constructs supplemented with 100 μM ZnCl2), followed by induction with 0.4 mM IPTG. Cells were left overnight at 18 °C, shaking 210 rpm. Cells were pelleted, flash frozen, and stored at -80 °C.

### Protein Purification

GST-tagged full-length H/K/NRAS (HRAS 1-189; KRAS 1-188; NRAS 1-189) pellet was thawed and resuspended in lysis buffer (20 mM HEPES pH 7.4, 150 mM NaCl, 1 mM EDTA, 5% Glycerol, and protease inhibitor cocktail). Whole cell lysate was passed through the French pressure cell press at 1520 psi and centrifuged. Soluble cell lysate was incubated with pre-equilibrated glutathione resin for 1 hour at 4°C. After extensive washing, protein was eluted off the resin with elution buffer (20 mM HEPES pH 7.4, 150 mM NaCl, 1 mM EDTA, 5% Glycerol, and 10 mM reduced glutathione). To dissociate bound nucleotide, proteins were incubated in HEPES buffer with 10 mM EDTA and 10-fold molar excess of GMPPNP (Sigma-Aldrich) for 30 min at 4 °C. To allow rebinding, MgCl2 was added to a final concentration of 1 mM and rotated for 2 hours at 4 °C. HRAS/KRAS were further purified on a Superdex 200 10/300 GL size exclusion chromatography column (Cytiva). Main steps were checked with SDS-PAGE followed by Coomassie staining. After concentration, aliquots were flash frozen and stored at - 80°C. No post translational modifications were added to RAS proteins.

C/ARAF pellets were thawed and resuspended in lysis buffer (20 mM HEPES pH 7.4, 150 mM NaCl, 5% Glycerol, 5mM TCEP, and protease inhibitor cocktail). Whole cell lysate was passed through the French pressure cell press at 1520 psi and centrifuged. Supernatant was incubated with pre-equilibrated Ni-NTA resin for 1 h at 4°C. After extensive washing, protein was eluted off the resin with elution buffer (20 mM HEPES pH 7.4, 150 mM NaCl, 5% glycerol, and increasing imidazole concentrations [90, 200, 400 mM]). Protein was further purified on a Superdex 200 10/300 GL size exclusion chromatography column (Cytiva). Main steps were checked with SDS-PAGE followed by Coomassie staining. After concentration, aliquots were flash frozen and stored at - 80°C.

### Pulldowns

Equimolar amounts of each protein were combined and allowed to associate in binding buffer (20 mM HEPES pH 7.4, 150 mM NaCl, 5% glycerol, 0.125 mg/mL BSA). The mixture was then incubated with 20 µL of Glutathione Sepharose 4B (Cytiva) to capture tagged proteins and their interacting partners. Resin-bound complexes were washed extensively with a high-salt buffer (20 mM HEPES pH 7.4, 500 mM NaCl, 5% glycerol, 0.125 mg/mL BSA). Following the final wash, 30 µL of 4x loading dye was added directly to the resin, and samples were analyzed by SDS–PAGE. Proteins were transferred to nitrocellulose membranes and detected using the following primary antibodies: anti-GST for GST-HRAS, GST-KRAS, and GST-NRAS (Santa Cruz Biotechnology, SC-138) and anti-His for His-tagged A/CRAF N-terminal fragments or CRAF kinase domain (Sigma, SAB5600227). Immunoblots were imaged using a LICOR scanner.

### Biolayer Interferometry (BLI)

Binding interactions between RAF constructs and RAS proteins were measured using an Octet N1 biolayer interferometry instrument (Sartorius). His-tagged ARAF or CRAF constructs were immobilized onto Anti-Penta-HIS (HIS1K) biosensors by loading 4 μL of protein at concentrations of 10-35 ng/μL to achieve an immobilization response of ∼1.0-1.2 nm. All experiments were performed in buffer containing 20 mM HEPES (pH 7.4), 150 mM NaCl, 0.05% Tween-20, with glycerol concentrations matched to the corresponding RAS protein preparations. Biosensor loading was performed for 3-5 min at 2000 rpm shaking speed, followed by equilibration in buffer for 1-2 min until a stable baseline was established.

RAS analytes (HRAS, KRAS, NRAS, or CRAF kinase domain where indicated) were prepared in the same buffer and tested over a concentration range of 12.4 nM to 20 μM. Association was monitored for 1-3 min by dipping loaded sensors into 4 μL analyte solution, followed by dissociation in 400 μL assay buffer for 1-5 min at 2000 rpm shaking speed. For inhibitor-binding experiments, full-length NRAS or KRAS4b was loaded with GMPPNP to model the active GTP-bound state prior to incubation with inhibitor and BLI analysis. Binding curves were globally fit using a 1:1 binding model in Octet N1.4 analysis software to determine kinetic parameters and equilibrium dissociation constants (*K_D_*). Data visualization and curve plotting were performed using GraphPad Prism.

All quantitative measurements represent at least two independent biological replicates performed using the same purified protein preparations. Reported *K_D_* values represent the mean of all replicate measurements, and error values correspond to the standard deviation (SD) across replicates.

### HDX-MS

CRAF RBD (102 µM), CRAF RBD CRD (138 µM), and NRAS (27 µM) were prepared in 20 mM HEPES pH 7.4, 150 mM NaCl, and 5% glycerol. Deuterium exchange was initiated by diluting each protein 1:5 (vol/vol) into a matching D₂O buffer and allowed to proceed for time points ranging from 20 s to 4.5 h. Reactions were quenched 1:1 with ice-cold buffer (100 mM phosphate pH 2.4, 0.5 M TCEP, 3 M guanidinium chloride) and immediately digested on an immobilized pepsin column.

Peptides were trapped (Higgins TARGA C8, 5 µm, 5 x 1.0 mm), desalted at 0 °C, separated by reversed-phase chromatography (TARGA C8, 50 x 0.3 mm; 10–40% acetonitrile gradient over 15 min, 8 µL/min), and analyzed on a Thermo Q-Exactive mass spectrometer. Peptides were identified using MS/MS of undeuterated samples and searched with SEQUEST against a database containing CRAF RBD, CRAF RBD CRD, HRAS, KRAS, NRAS, pepsin, and common contaminants. A subset of reproducibly detected peptides (119 for CRAF RBD and 86 for CRAF RBD CRD) was used for HDX analysis.

Isotopic envelopes, deuterium incorporation values, and peptide coverage maps were generated using ExMS2 (Kan et al., 2019). All deuterium-uptake plots and downstream analyses were produced using custom Python scripts that processed ExMS2 output tables.

### NanoBiT Constructs Cloning

NanoBiT® CMV MCS BiBiT vector, which contains a BRAF N- terminal fused with LgBiT and CRAF N-terminal fused with SmBiT, were purchased from Promega.

KRAS^LgBiT^ and CRAF^SmBiT^ was generated using the standard Gibson Assembly (NEB HiFi Assembly) using NanoBiT® BiBiT as the vector and following primers: 5’ CTCCGCCCCCCAGCGACACCATAATTACACACTTTGTCTTTG 3’, 5’ CCACAAGTTTATATTCAGTCATGGCGATCGCTAGCTGCAAA 3’, 5’ GGCGATCGCTAGCTGCAAAAAGAACAAG 3’, 5’ TCCGCCCCCCAGCGACAC 3’, and NRAS^LgBiT^ and CRAF^SmBiT^ was generated using the standard Gibson Assembly using NanoBiT® BiBiT as the vector and following primers: 5’ CCTCCGCCCCCCAGCGACACCATCACCACACATGGCAATC 3’, 5’ CACCAGTTTGTACTCAGTCATGGCGATCGCTAGCTGCAAAA 3’, 5’ GGCGATCGCTAGCTGCAAAAAGAACAAG 3’, 5’ TCCGCCCCCCAGCGACAC 3’.

Primers were designed to truncate CRAF^SmBiT^ to only include the RBD and CRD regions for NanoBiT. Primers were designed to introduce the desired truncation. The forward primer sequences were 5’ ATGTCTAAGACAAGCAACACTATCCG 3’ and 5’ GTTTCTCTCGGAGGAGGT 3’ and the reverse primer sequences were 5’ ATAGGGCTAGCGATCGCC 3’ and 5’ TATGTGTGTGGACTGGAGT 3’. Standard Gibson Assembly (NEB HiFi Assembly) was carried out using NanoBiT® BiBiT as the vector.

### NanoBiT Luminescence Assay

HEK293 cells were transiently transfected with wild-type full-length (WTFL) CRAF^SmBiT^-KRAS^LgBiT^, WTFL CRAF^SmBiT^-NRAS^LgBiT^, RBD CRD CRAF^SmBiT^-KRAS^LgBiT^, or RBD CRD CRAF^SmBiT^-NRAS^LgBiT^ at 2 µg plasmid per well and incubated at 37°C. Twenty-four hours post-transfection, cells were trypsinized, pelleted, resuspended in OptiMEM+4% FBS, and re-seeded onto 96-well plate at 5×10^4^ cells per well then incubated at 37°C overnight. Cells were treated with 10 µM of Sotorasib, Adagrasib, BI 2865, or RMC 6236 for 4 hours. After treatment, 10 µM furimazine substrate was added to each well and luminescence readings were taken at 470-480 nm using CLARIOstar plate reader. Statistical significance was determined through paired t test where indicated, followed by Tukey’s honest significant difference post-hoc test with corresponding P-values (*P<0.05, **P<0.01, ***P<0.001). All graph bars represent the mean ± SD with individual data points per biological replicate with corresponding p-values.

**Table 1.** Antibodies used in immunoblots. **Inhibitors.** BI-2865 (E1474), RMC-6236 (E1597), Sotorasib (S8830), and Adagrasib (S8884) were purchased from Selleckchem.

| **Manufacturer** | **Catalog Number** | **Name** |
| --- | --- | --- |
| VWR | 89356-446 | GST |
| Santa Cruz | sc-8036 | His |
| Cell Signaling | 9422 | CRAF |
| Cell Signaling | 9421 | CRAF pS259 |
| Cell Signaling | 67648 | RAS |
| Cell Signaling | 3700 | β-Actin |
| Licor | 926-32211 | Anti-rabbit |
| Licor | 926-68070 | Anti-mouse |

## Declaration of Interests

The authors declare no competing interests.

## Funding Sources

The authors declare their funding sources as follows: Rowan University Department of Chemistry and Biochemistry Fellowship to SB, WW Smith Charitable Trust to ZW, and NIH funding R01GM138671 to ZW.

## Data Sharing Plan

All data are included in the manuscript and supporting information.

## Notes

### Competing Interest Statement

The authors have declared no competing interest.

