## Supplementary Files for "Divergent CRD-Dependent Mechanisms Govern RAS Isoform-Selective Recruitment of CRAF and ARAF"

### Supplementary Information

- Supplementary Figures 1-7
- Supplementary Table 1 and 2

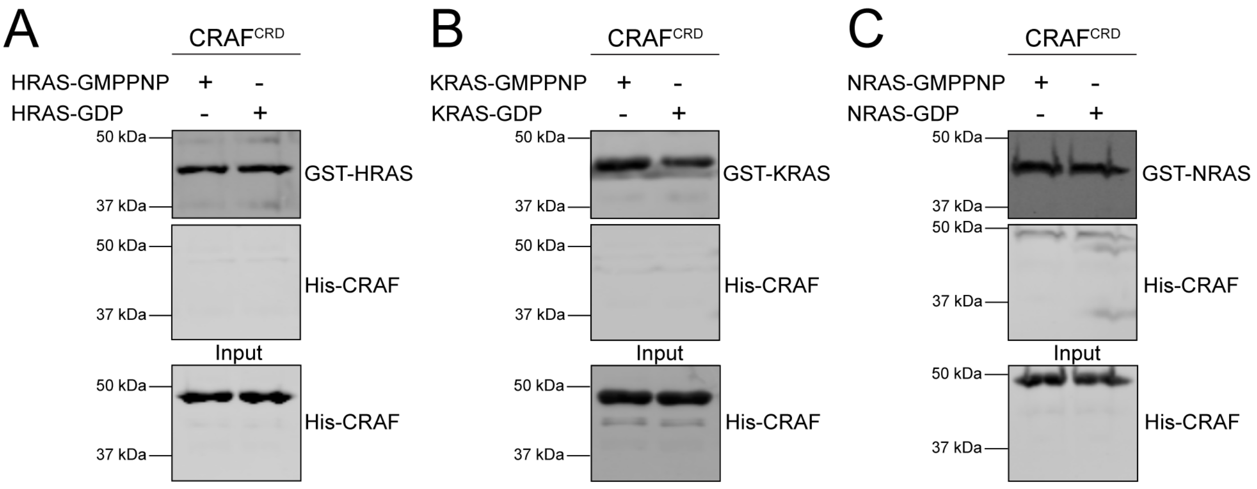


**Figure S1. The isolated CRAF$^{CRD}$ is insufficient for stable interaction with RAS isoforms.** A-C) GST pulldown assays assessing binding of the isolated His-tagged CRAF$^{CRD}$ to HRAS (A), KRAS (B), or NRAS (C) loaded with either GMPPNP or GDP. Pulldown experiments were performed in two independent replicates with comparable results.

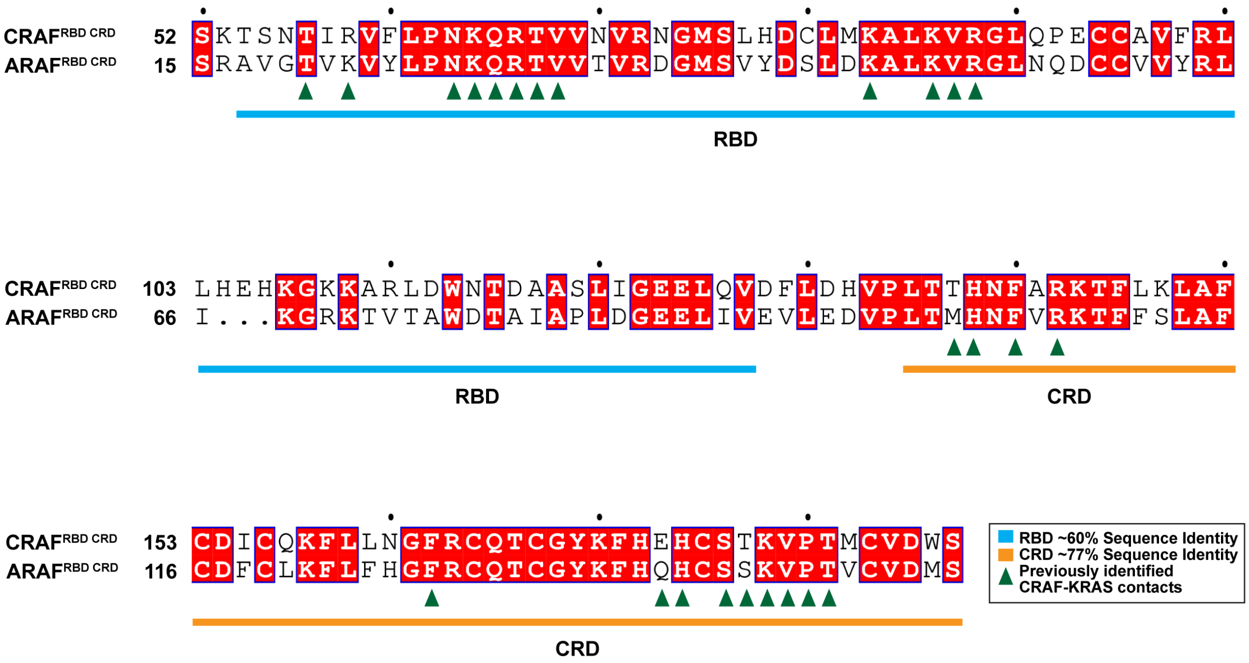


**Figure S2. Sequence alignment of CRAF and ARAF RBD and CRD domains highlighting conservation and RAS-interacting residues.** Pairwise sequence alignment of the RBD and CRD of human CRAF and ARAF. Residues are colored according to sequence identity (red shading), with identical residues highlighted. Domain boundaries for the RBD (blue) and CRD (orange) are indicated below the alignment. CRAF residues previously identified to participate in KRAS interaction are marked (green triangles).

A

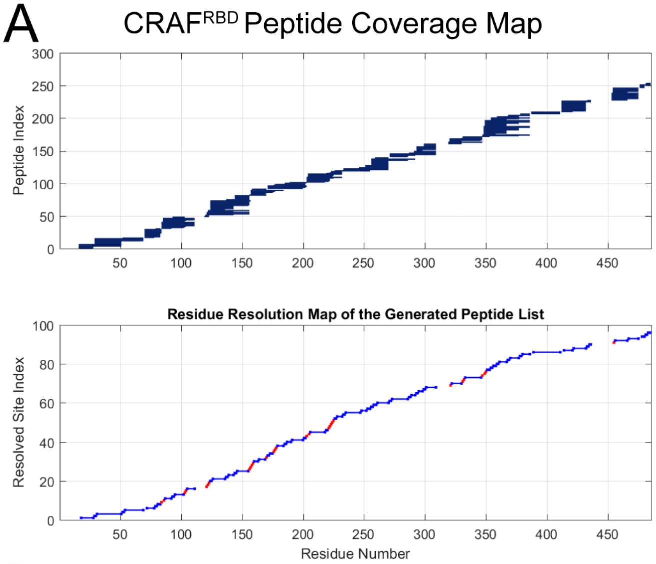


B

CRAF$^{RBD}$ Sequence:
1- HHHHHHGTKTEEGKLVIWINGDKGYNGLAEVGKKFEKDTGIKVTVEHPDKL
EEKFPQVAATGDGPDIIFWAHDRFGGYAQSGLLAEITPDKAFQDKLYPFTWDA
VRYNGKLIAYPIAVEALSLIYNKDLLPNPPKTWEEIPALDKELKAKGKSALMFNL
QEPYFTWPLIAADGGYAFKYENGKYDIKDVGVDNAGAKAGLTFLVDLIKNKHM
NADTDYSIAEAAFNKGETAMTINGPWAWSNIDTSKVNYGVTVLPTFKGQPSKP
FVGVLSAGINAASPNKELAKEFLENYLLTDEGLEAVNKDKPLGAVALKSYEEEL
AKDPRIAATMENAQKGEIMPNIPQMSAFWYAVRTAVINAASGRQTVDEALKDA
QTGTDYDIPTTENLYFQGHMASMTGGQQMGRGSSKTSNTIRVFLPNKQRTV
VNVRNGMSLHDCLMKALKVRGLQPECCAVFRLLHEHKGKKARLDWNTDAAS
LIGEELQVDFL -485

His/MBP tag
CRAF$^{RBD}$ (a.a. 52-131)

**Figure S3. Peptide coverage and sequence representation of the CRAF$^{RBD}$ used for HDX-MS analysis.** A) Stripe plots depicting peptide coverage across the CRAF$^{RBD}$, showing identified peptides (top) and the corresponding residue resolution map generated from the peptide list (bottom). B) Amino acid sequence of the CRAF$^{RBD}$ construct used for HDX-MS, with the region analyzed highlighted in purple and the N-terminal His/MBP tag indicated in green. Residue numbering corresponds to the full-length CRAF sequence.

A

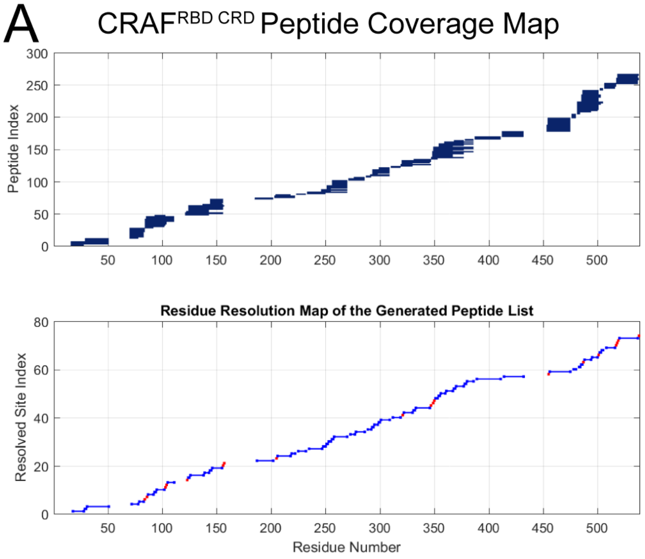


B

CRAF$^{RBD\ CRD}$ Sequence:
1- HHHHHHGTKTEEGKLVIWINGDKGYNGLAEVGKKFEKDTGIKVTVEHPDKL
EEKFPQVAATGDGPDIIFWAHDRFGGYAQSGLLAEITPDKAFQDKLYPFTWDA
VRYNGKLIAYPIAVEALSLIYNKDLLPNPPKTWEEIPALDKELKAKGKSALMFNL
QEPYFTWPLIAADGGYAFKYENGKYDIKDVGVDNAGAKAGLTFLVDLIKNKHM
NADTDYSIAEAAFNKGETAMTINGPWAWSNIDTSKVNYGVTVLPTFKGQPSK
PFVGVLSAGINAASPNKELAKEFLENYLLTDEGLEAVNKDKPLGAVALKSYEEE
LAKDPRIAATMENAQKGEIMPNIPQMSAFWYAVRTAVINAASGRQTVDEALKD
AQTGTDYDIPTTENLYFQGHMASMTGGQQMGRGSSKTSNTIRVFLPNKQRT
VVNVRNGMSLHDCLMKALKVRGLQPECCAVFRLLHEHKGKKARLDWNTDA
ASLIGEELQVDFLDHVPLTTHNFARKTFLKLAFCDICQKFLLNGFRCQTCGYKF
HEHCSTKVPTMCVDWS -542

His/MBP tag
CRAF$^{RBD\ CRD}$ (a.a. 52-188)

**Figure S4. Peptide coverage and sequence representation of the CRAF$^{RBD\ CRD}$ used for HDX-MS analysis.** A) Stripe plots depicting peptide coverage across the CRAF$^{RBD\ CRD}$, showing identified peptides (top) and the corresponding residue resolution map generated from the peptide list (bottom). B) Amino acid sequence of the CRAF$^{RBD\ CRD}$ construct used for HDX-MS, with the region analyzed highlighted in brown and the N-terminal His/MBP tag indicated in green. Residue numbering corresponds to the full-length CRAF sequence.

A

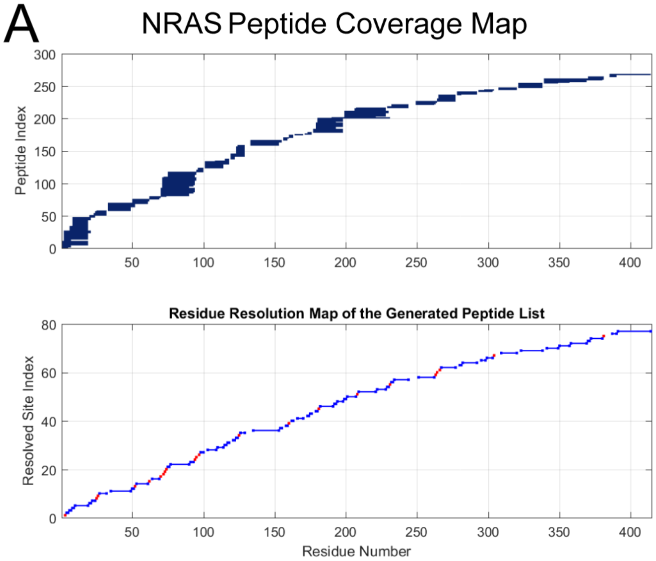


B

NRAS Sequence:
1- MSPILGYWKIKGLVQPTRLLLEYLEEKYEEHLYERDEGDKWRNKKFELGLE
FPNLPYYIDGDVKLTQSMAIIRYIADKHNMLGGCPKERAEISMLEGAVLDIRYG
VSRIAYSKDFETLKVDFLSKLPEMLKMFEDRLCHKTYLNGDHVTHPDFMLYDA
LDVVLYMDPMCLDAFPKLVCFKKRIEAIPQIDKYLKSSKYIAWPLQGWQATFGG
GDHPPKSDLVPRGSMTEYKLVVVGAGGVGKSALTIQLIQNHFVDEYDPTIEDSY
RKQVVIDGETCLLDILDTAGQEEYSAMRDQYMRTGEGFLCVFAINNSKSFADI
NLYREQIKRVKDSDDVPMVLVGNKCDLPTRTVDTKQAHELAKSYGIPFIETSAK
TRQGVEDAFYTLVREIRQYRMKKLNSSDDGTQGCMGLPCVVM -414

GST tag
NRAS (a.a. 1-189)

**Figure S5. Peptide coverage and sequence representation of NRAS used for HDX-MS analysis.** A) Stripe plots depicting peptide coverage across NRAS, showing identified peptides (top) and the corresponding residue resolution map generated from the peptide list (bottom). B) Amino acid sequence of NRAS construct used for HDX-MS, with the region analyzed highlighted in pink and the N-terminal GST tag indicated in green.

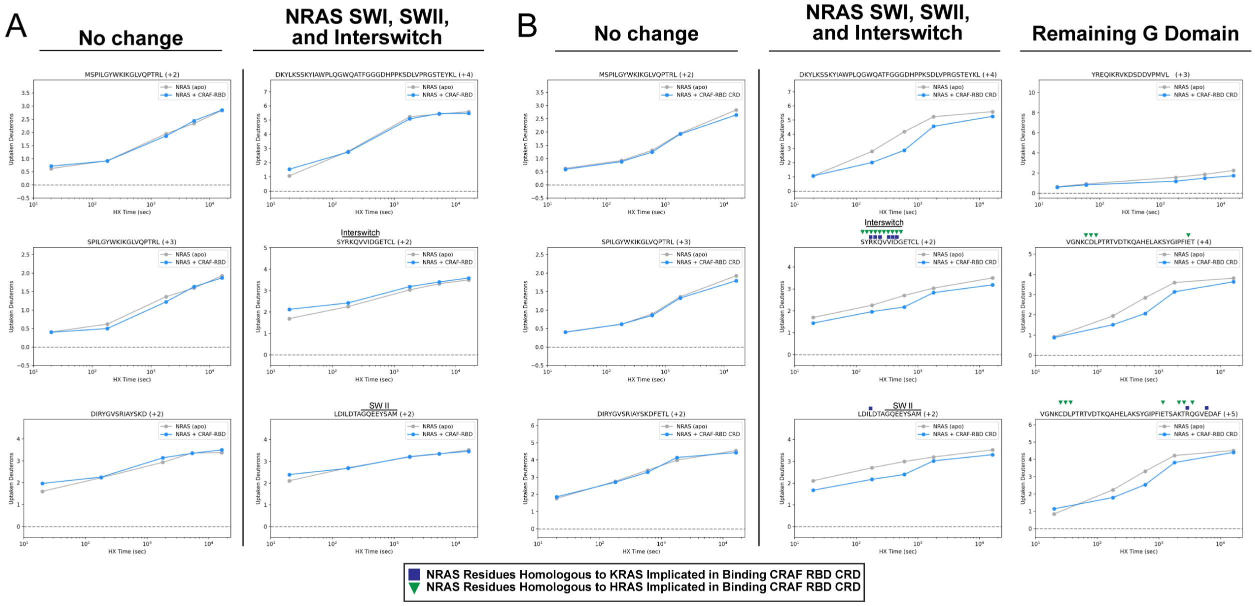


**Figure S6. Reciprocal HDX-MS analysis comparing apo NRAS with NRAS bound to CRAF$^{RBD}$ or CRAF$^{RBD\ CRD}$.** A) Representative deuterium uptake plots for NRAS peptides exhibiting no significant change in exchange in the presence of CRAF$^{RBD}$. B) Deuterium uptake plots for peptides spanning NRAS SWI, SWII, interswitch regions, and additional regions within the remaining G-domain that display reduced deuterium uptake in the presence of CRAF$^{RBD\ CRD}$, resulting in reproducible reductions in deuterium incorporation consistent with stabilization of the effector-binding interface. NRAS residues homologous to KRAS (blue squares) and HRAS (green triangle) residues previously implicated in binding CRAF$^{RBD\ CRD}$ have been indicated.

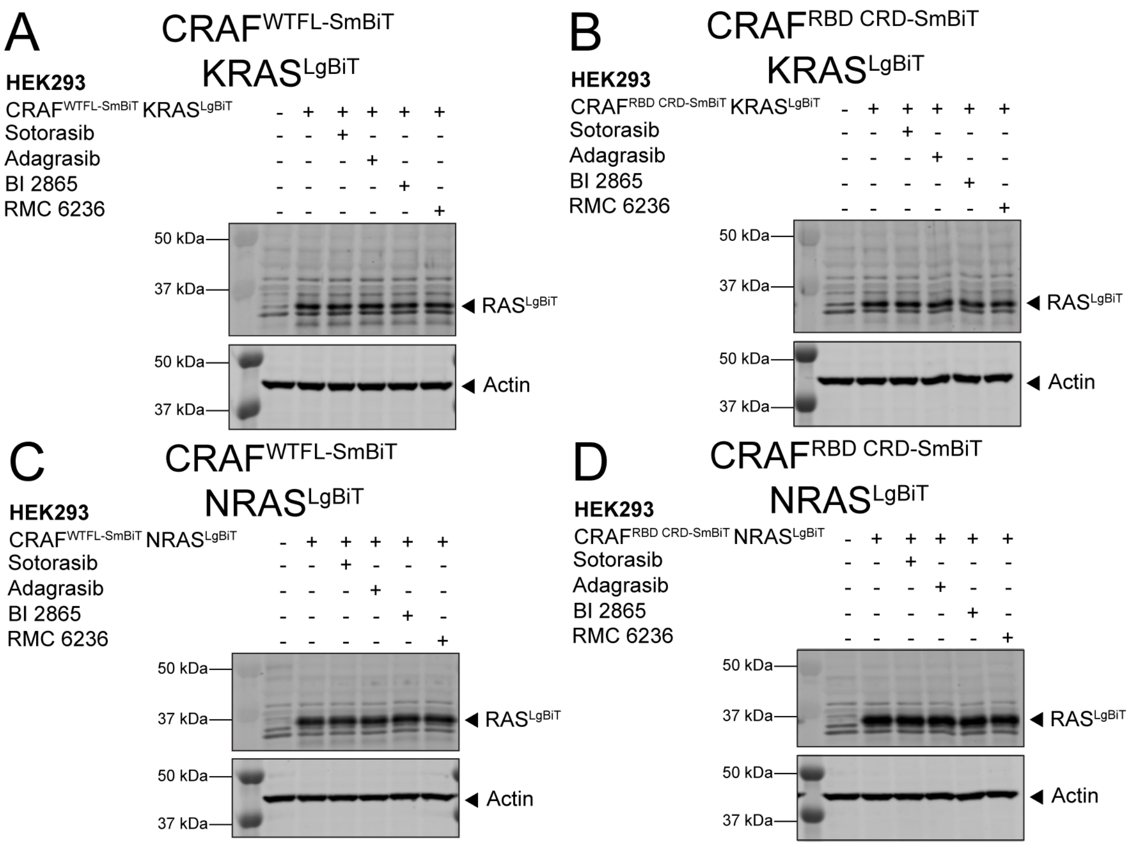


**Figure S7. RAS inhibitors disrupt RAS–CRAF interactions without altering protein expression in cells.** A-B) Representative immunoblots of HEK293 cells expressing $CRAF^{WTFL\text{-}SmBiT}$ with $KRAS^{LgBiT}$ (A) or $CRAF^{RBD\ CRD\text{-}SmBiT}$ with $KRAS^{LgBiT}$ (B) in the absence or presence of the indicated RAS inhibitors (Sotorasib, Adagrasib, BI 2865, and RMC 6236). C-D) Immunoblots of HEK293 cells expressing $CRAF^{WTFL\text{-}SmBiT}$ with $NRAS^{LgBiT}$ (C) or $CRAF^{RBD\text{-}CRD\text{-}SmBiT}$ with $NRAS^{LgBiT}$ (D) under the same treatment conditions.

#### Table S1. BLI Kinetic Data Summary

| Binding Partners | $K_D$ (nM) | $K_{on}$ (1/Ms) | $K_{off}$ (1/s) | $X^2$ | $R^2$ |
|---|---|---|---|---|---|
| $CRAF^{RBD}$-HRAS | 63.23 | $1.155 \times 10^4$ | $7.303 \times 10^{-4}$ | 3.65 | 0.993 |
| | 40.24 | $2.455 \times 10^4$ | $9.880 \times 10^{-4}$ | 0.43 | 0.997 |
| $CRAF^{RBD}$-KRAS | 96.66 | $3.107 \times 10^4$ | $3.004 \times 10^{-3}$ | 1.90 | 0.997 |
| | 101.77 | $4.926 \times 10^4$ | $5.012 \times 10^{-3}$ | 3.46 | 0.992 |
| $CRAF^{RBD}$-NRAS | 325.7 | $1.137 \times 10^4$ | $3.704 \times 10^{-3}$ | 1.64 | 0.987 |
| | 398.0 | $1.154 \times 10^4$ | $4.595 \times 10^{-3}$ | 1.54 | 0.990 |
| $CRAF^{RBD\ CRD}$-HRAS | 61.15 | $2.458 \times 10^4$ | $1.503 \times 10^{-3}$ | 0.70 | 0.987 |
| | 51.17 | $6.272 \times 10^4$ | $3.210 \times 10^{-3}$ | 0.94 | 0.994 |
| $CRAF^{RBD\ CRD}$-KRAS | 30.31 | $1.040 \times 10^3$ | $3.154 \times 10^{-5}$ | 3.11 | 0.992 |
| | 34.80 | $1.390 \times 10^3$ | $4.838 \times 10^{-5}$ | 2.74 | 0.971 |
| $CRAF^{RBD\ CRD}$-NRAS | 73.41 | $2.452 \times 10^4$ | $1.800 \times 10^{-3}$ | 0.51 | 0.994 |
| | 74.88 | $6.832 \times 10^4$ | $5.116 \times 10^{-3}$ | 1.14 | 0.989 |
| $ARAF^{RBD}$-HRAS | 2493 | $9.507 \times 10^2$ | $2.370 \times 10^{-3}$ | 0.66 | 0.981 |
| | 6838 | $3.684 \times 10^2$ | $2.519 \times 10^{-3}$ | 2.24 | 0.978 |
| $ARAF^{RBD}$-KRAS | 966.65 | $3.068 \times 10^2$ | $2.966 \times 10^{-4}$ | 0.69 | 0.971 |
| | 1519 | $2.950 \times 10^2$ | $4.481 \times 10^{-4}$ | 0.25 | 0.983 |
| $ARAF^{RBD}$-NRAS | 8289 | $1.007 \times 10^3$ | $1.575 \times 10^{-3}$ | 0.63 | 0.975 |
| | 6082 | $1.007 \times 10^3$ | $6.122 \times 10^{-3}$ | 1.60 | 0.984 |
| $ARAF^{RBD\ CRD}$-HRAS | 7294 | $4.866 \times 10^2$ | $3.550 \times 10^{-3}$ | 0.28 | 0.980 |
| | 11981 | $2.367 \times 10^2$ | $2.836 \times 10^{-3}$ | 0.92 | 0.918 |
| $ARAF^{RBD\ CRD}$-KRAS | 1888 | $9.619 \times 10^2$ | $1.816 \times 10^{-3}$ | 0.20 | 0.964 |
| | 4449 | $2.193 \times 10^2$ | $9.757 \times 10^{-4}$ | 1.49 | 0.945 |
| $ARAF^{RBD\ CRD}$-NRAS | 1227 | $4.291 \times 10^2$ | $5.267 \times 10^{-4}$ | 0.66 | 0.990 |
| | 4268 | $3.477 \times 10^2$ | $1.484 \times 10^{-3}$ | 2.45 | 0.978 |
| $CRAF^{RBD\ CRD}$-$CRAF^{KD}$ | 47.53 | $1.580 \times 10^6$ | $7.509 \times 10^{-2}$ | 0.02 | 0.994 |
| | 39.72 | $1.181 \times 10^6$ | $4.691 \times 10^{-2}$ | 0.10 | 0.976 |
| $CRAF^{RBD\ CRD}$+KRAS-$CRAF^{KD}$ | 600.16 | $1.295 \times 10^5$ | $7.772 \times 10^{-2}$ | 0.06 | 0.945 |
| | 519.91 | $1.377 \times 10^5$ | $7.160 \times 10^{-2}$ | 0.24 | 0.894 |
| $CRAF^{RBD\ CRD}$-KRAS+BI2865 | 713.77 | $1.114 \times 10^4$ | $7.953 \times 10^{-3}$ | 1.17 | 0.945 |
| | 421.87 | $7.406 \times 10^4$ | $2.145 \times 10^{-4}$ | 0.70 | 0.957 |
| $CRAF^{RBD\ CRD}$-KRAS+RMC6236 | 1901 | $3.986 \times 10^4$ | $7.577 \times 10^{-2}$ | 1.73 | 0.947 |
| | 1496 | $5.304 \times 10^4$ | $7.935 \times 10^{-2}$ | 0.38 | 0.983 |
| $CRAF^{RBD\ CRD}$-NRAS+RMC6236 | 633.54 | $6.617 \times 10^4$ | $4.192 \times 10^{-2}$ | 0.40 | 0.991 |
| | 614.45 | $5.554 \times 10^4$ | $3.413 \times 10^{-2}$ | 0.46 | 0.97 |

**Table S2. List of Previously Published $K_D$ Values**

| RAF Construct | RAS Construct | Binding Affinity (nM) | Method | Expression System | Reference |
|---|---|---|---|---|---|
| $CRAF^{RBD}$ (51-131) | HRAS | 270 ± 90 | Total internal reflection fluorescence (TIRF) microscopy (2% PS) | E.coli | (Jimenez Salinas et al., 2026) |
| $CRAF^{RBD}$ (51-131) | HRAS | 240 ± 20 | TIRF (20% PS) | E.coli | (Jimenez Salinas et al., 2026) |
| $CRAF^{RBD\ CRD}$ (51-188) | HRAS | 25 ± 10 | TIRF (2% PS) | E.coli | (Jimenez Salinas et al., 2026) |
| $CRAF^{RBD\ CRD}$ (51-188) | HRAS | 6 ± 3 | TIRF (20% PS) | E.coli | (Jimenez Salinas et al., 2026) |
| $CRAF^{CRD}$ (138-188) | HRAS | NB | TIRF (20% PS) | E.coli | (Jimenez Salinas et al., 2026) |
| $CRAF^{RBD}$ (51-131) | HRAS | 220 ± 80 | TIRF (PEGylated) | E.coli | (Jimenez Salinas et al., 2026) |
| $CRAF^{RBD\ CRD}$ (51-188) | HRAS | 40 ± 10 | TIRF (PEGylated) | E.coli | (Jimenez Salinas et al., 2026) |
| $CRAF^{RBD}$ (52-131) | KRAS (1-169) | 356 | Surface Plasmon Resonance (SPR) | E.coli | (Tran et al., 2021) |
| $CRAF^{RBD\ CRD}$ (52-192) | KRAS (1-169) | 152 | SPR | E.coli | (Tran et al., 2021) |
| $CRAF^{RBD}$ (51-131) | KRAS (1-166) | 98 ± 2 | Isothermal Titration Calorimetry (ITC) | E.coli | (Johnson et al., 2017) |

| | | | | | |
|---|---|---|---|---|---|
| $CRAF^{RBD}$ (51-131) | HRAS (1-166) | 94 ± 4 | ITC | E.coli | (Johnson et al., 2017) |
| $CRAF^{RBD}$ (51-131) | NRAS (1-166) | 206 ± 8 | ITC | E.coli | (Johnson et al., 2017) |
| $CRAF^{RBD}$ (1-149) | HRAS (farnesylated) | 24.3 | SPR | E.coli | (Fischer et al., 2007) |
| $CRAF^{RBD}$ (1-149) | $HRAS^{G12V}$ (farnesylated) | 21.3 | SPR | E.coli | (Fischer et al., 2007) |
| $CRAF^{RBD}$ (1-149) | HRAS | 21.2 | SPR | E.coli | (Fischer et al., 2007) |
| $BRAF^{RBD}$ (1-245) | HRAS | 11.2 | SPR | E.coli | (Fischer et al., 2007) |
| $ARAF^{RBD}$ | HRAS | 76.6 | SPR | E.coli | (Fischer et al., 2007) |
| $CRAF^{FL}$ | HRAS (farnesylated) | 460 | SPR | HRAS: E.coli CRAF: Sf9 | (Fischer et al., 2007) |
| $CRAF^{R89L}$ | HRAS (farnesylated) | 860 | SPR | HRAS: E.coli CRAF: Sf9 | (Fischer et al., 2007) |
| $CRAF^{FL}$ | HRAS | NB | SPR | HRAS: E.coli CRAF: Sf9 | (Fischer et al., 2007) |
| $BRAF^{FL}$ | HRAS (farnesylated) | 58.8 | SPR | HRAS: E.coli BRAF: Sf9 | (Fischer et al., 2007) |
| $BRAF^{FL}$ | HRAS | 80.2 | SPR | HRAS: E.coli BRAF: Sf9 | (Fischer et al., 2007) |
| $BRAF^{\Delta 98}$ (99-766) | HRAS (farnesylated) | 75.6 | SPR | HRAS: E.coli BRAF: Sf9 | (Fischer et al., 2007) |
| $BRAF^{\Delta 98}$ (99-766) | HRAS | 290.4 | SPR | HRAS: E.coli BRAF: Sf9 | (Fischer et al., 2007) |
| $CRAF^{RBD}$ CRD (1-192) | HRAS (farnesylated) | 122.3 | SPR | E.coli | (Fischer et al., 2007) |
| $CRAF^{RBD}$ CRD | HRAS | 190.5 | SPR | E.coli | (Fischer et al., 2007) |

| | | | | | |
|---|---|---|---|---|---|
| (1-192) | | | | | |
| $CRAF^{N-terminal}$ (1-303) | HRAS (farnesylated) | 104.6 | SPR | E.coli | (Fischer et al., 2007) |
| $CRAF^{N-terminal}$ (1-303) | HRAS | 188 | SPR | E.coli | (Fischer et al., 2007) |
| $BRAF^{N-terminal}$ (1-412) | HRAS (farnesylated) | 62.4 | SPR | E.coli | (Fischer et al., 2007) |
| $BRAF^{N-terminal}$ (1-412) | HRAS | 64.4 | SPR | E.coli | (Fischer et al., 2007) |
| $CRAF^{RBD}$ (51-131) | Ras·mantGppNHp | 160 | SPR | E.coli | (Sydor et al., 1998) |
| $CRAF^{RBD}_{L91W/H}$ (51-131) | H + Ras·mantGppNHp | 780 | SPR | E.coli | (Sydor et al., 1998) |
| $CRAF^{RBD}$ (51-131) | Ras·mantGppNHp | 300 | SPR | E.coli | (Sydor et al., 1998) |
| $CRAF^{RBD}_{L91W/H}$ (51-131) | H + Ras·GppNHp | 380 | SPR | E.coli | (Sydor et al., 1998) |
| $CRAF^{RBD}$ (51-131) | Ras·mantGDP | 32,000 | SPR | E.coli | (Sydor et al., 1998) |
